# Genetic differences between extreme and composite constitution types from whole exome sequences reveal actionable variations

**DOI:** 10.1101/2020.04.24.059006

**Authors:** Tahseen Abbas, Rintu Kutum, Rajesh Pandey, Pushkar Dakle, Ankita Narang, Vijeta Manchanda, Rutuja Patil, Dheeraj Aggarwal, Gourja Bansal, Pooja Sharma, Gaura Chaturvedi, Bhushan Girase, Ankita Srivastava, Sanjay Juvekar, Debasis Dash, Bhavana Prasher, Mitali Mukerji

## Abstract

Personalized medicine relies on successful identification of genome-wide variations that governs inter-individual differences in phenotypes and system level outcomes. In Ayurveda, assessment of composite constitution types “*Prakriti”* forms the basis for risk stratification, predicting health and disease trajectories and personalized recommendations. Here, we report a novel method for identifying pleiotropic genes and variants that associate with healthy individuals of three extreme and contrasting “*Prakriti”* constitutions through exome sequencing and state-of-the-art computational methods. Exome Seq of three extreme *Prakriti* types from 108 healthy individuals 54 each from genetically homogeneous populations of North India (NI, Discovery cohort) and Western India (VADU, Replication cohort) were evaluated. Fisher’s Exact Test was applied between *Prakriti* types in both cohorts and further permutation based p-value was used for selection of exonic variants. To investigate the effect of sample size per genetic association test, we performed power analysis. Functional impact of differentiating genes and variations were inferred using diverse resources -Toppfun, GTEx, GWAS, PheWAS, UK Biobank and mouse knockdown/knockout phenotype (MGI). We also applied supervised machine learning approach to evaluate the association of exonic variants with multisystem phenotypes of *Prakriti*. Our targeted investigation into exome sequencing from NI (discovery) and VADU (validation) cohorts datasets provide ~7,000 differentiating SNPs. Closer inspection further identified a subset of SNPs (2407 (NI) and 2393 (VADU)), that mapped to an overlapping set of 1181 genes. This set can robustly stratify the Prakriti groups into three distinct clusters with distinct gene ontological (GO) enrichments. Functional analysis further strengthens the potential pleiotropic effects of these differentiating genes/variants and multisystem phenotypic consequences. Replicated SNPs map to some very prominent genes like *FIG4, EDNRA, ANKLE1, BCKDHA, ATP5SL, EXOCS5*, *IFIT5, ZNF502, PNPLA3 and IL6R*. Lastly, multivariate analysis using random forest uncovered rs7244213 within urea transporter *SLC14A2*, that associate with an ensemble of features linked to distinct constitutions. Our results reinforce the concept of integration of Prakriti based deep phenotypes for risk stratification of healthy individuals and provides markers for early actionable interventions.

## Background

Precision medicine aims to stratify individuals based on endo-phenotypes and risk profiles, for early actionable interventions [1]. Methods are being evolved to identify biomarkers corresponding to phenotypes that could enable screening of target populations, predict progression and prognosis of illness, as well as enable differential therapeutic managements [1–5]. Successful mapping of an individual’s genotype to phenotype and health trajectories in order to predict systemic outcomes is a challenge, as our understanding of the human phenomic architecture from genomic profiles is still in its infancy. Most of the Genome-Wide Association Studies (GWAS) for delineation of the genetic basis of common and complex diseases have been conducted on discernible traits; in the absence of comprehensive deeper phenotypes of multisystem attributes, much of the phenotype to genotype associations still remains to be uncovered [6]. Widespread overlap of GWAS SNP’s association with seemingly unrelated diseases and phenotypes [7,8] have prompted Phenome Wide Association Studies (PheWAS) in Biobanks, Electronic Health Records (EHR) as well as longitudinal cohorts [9–13]. PheWAS has uncovered many variants that exhibit pleiotropic effects and offers to identify disease gene networks, novel phenotypic associations of drugs side effects and leads for drug repurposing [14–17]. The success in uncovering phenotype-phenotype connectivity in PheWAS depends on the extent and diversity of captured features, as well as, the co-occurrence of the phenotypes in the EHRs and cohorts [18]. Thus even if the cohort size in PheWAS might be in millions with significant GWAS associations, the subsequent genotype-phenotype associations of the variants are in relatively smaller sample sizes. It is being felt that extending the GWAS to systems’ level with deeper phenotypes and composite traits can accelerate predictive marker discoveries [6,19]. Exome sequencing of extreme phenotypes in smaller sample sizes (i.e. hundreds) is also being used as another approach to identify variants with larger phenotypic effects in single attributes or for variable outcomes [20–24].

Ayurveda, the oldest documented system of personalized medicine, provides a rich repertoire of phenotypic descriptions for a comprehensive assessment of an individual’s constitution *Prakriti* types [25,26].We have previously organized these descriptions into a questionnaire of ~150 attributes to predict an individual *Prakriti* type [27,28]. Knowledge of *Prakriti* is fundamental in Ayurveda for the prediction of an individual health and disease trajectory [25], as well as for personalized management and therapy [27,28]. The three basic *Prakriti* types are Vata (V), Pitta (P) and Kapha (K), which give rise to seven constitution types (V, P, K, VP, VK, KP, VKP). The three basic *Prakriti* types (V, P or K only) comprise nearly 10% of the population [25].

They display extreme phenotypic variations, with highly contrasting drug responses and disease susceptibilities [25,27]. Unsupervised machine learning and advanced statistical approaches on phenotypes of healthy individuals of extreme *Prakriti* types have been used to validate the existence of *Prakriti* specific phenotype-phenotype connectivity [29]. Studies have also provided evidence for differences at different hierarchies of genetic, epigenetic, biochemical and microbiome between *Prakriti* types [28–33]. Using this approach, we have previously identified predictive markers in *EGLN1*, a gene linked to high altitude adaptation [32,34]. Allelic association of *EGLN1* to the hemostasis related platelet glycoprotein, von Willebrand factor (VWF) could explain thrombotic outcomes in high altitude hypoxic conditions for specific *Prakriti* types [32,34]. These studies effectively support the application of using the *Prakriti* phenotype scaffolds to identify key genetic features that govern systemic outcomes. Eventually, these leads serve as important biomarkers for pre-screening individuals prior to exposure, and also for targeted interventions.

We hypothesize that exome sequencing of healthy individuals of basic *Prakriti* types could enable discovery of variants with pleiotropic and penetrant effects. Here, we report the results from an in-depth exome sequencing analysis of 108 healthy individuals of basic constitution types across two genetically homogenous cohorts. Our analysis reveal significant exome-wide genetic differences between the *Prakriti* types, which are largely in line with the earlier reports. Importantly, we identified a set of 1181 overlapping genes that can robustly segregate the *Prakriti* types into three distinct clusters, in both the cohorts. Notably, a larger proportion of these differentiating variants are already reported in GWAS and PheWAS including the UK Biobank. Lastly, by multivariate analysis using random forest, we also demonstrate how some of the differentiating genotypes can predict an ensemble of phenotypes that distinguish *Prakriti*. This study provides a unique framework for enriching genetic markers associated with composite phenotypes that may be utilized for effective risk stratification of healthy individuals.

## Results

### Similar patterns of exonic differences amongst *Prakriti* types with significant overlap of genes between the cohorts

Recent reports suggest the utilization of exome sequencing in extreme phenotypes with unique attributes, as an alternative approach for the identification of highly penetrant genetic variants. We utilized this approach to comprehensively understand the underlying distribution of genetic variants in individuals with extreme composite and contrasting *Prakriti* types, from two genetically homogeneous cohorts (NI and VADU). A thorough investigation of this exome sequencing datasets revealed a total of 2,14,844 and 2,20,598 variants in NI and VADU cohorts, respectively (refer to the material and methods section). Comparative analysis assessing the distribution of the differentiating variants across the genic region revealed near similar distributions across both cohorts ruling out any bias in sequencing (Supplementary Fig.1, Additional File 2). Interestingly, about 50% of the differentiating variations map to exonic regions with a significant fraction in 3’UTRs (Supplementary Fig.1, Additional File 2). Further investigation of the variants that differ amongst *Prakriti* groups (V *vs* P, V *vs* K, and P *vs* K) led to the identification of 6534 unique variations (3749 genes) in NI and 7050 (3941 genes) in VADU (Supplementary Table 1, Additional File 1). Noteworthy, a total of 1181 genes with 472 identical SNPs were observed to be replicated between *Prakriti* groups across both cohorts (Supplementary Table 2, Additional File 1). Amongst these, 110 identical SNPs from 87 genes have similar profiles of frequency differences between *Prakriti* types e.g. in a V *vs* K comparison, a profile of V+K-represents higher alternate allele frequency in Vata compared to Kapha in both cohorts, despite the significant differences in frequency in the background population between the cohorts (Supplementary Table 1, Additional File 1).

### Population sub-stratification based on *Prakriti* differentiating SNPs

To evaluate if the identified *Prakriti* differentiating SNPs alone can sub-classify the population we first performed a Principal Component Analysis (PCA) with 2407 (NI) & 2393 (VADU) significant SNPs from 1181 replicated genes. The analysis revealed three distinct clusters for Vata, Pitta, and Kapha in both cohorts. Retaining the tag SNPs within the cohorts based on linkage disequilibrium (LD) still provided three distinct, albeit highly segregated clusters with a minimal set of 1605 (NI) SNPs and 1456 (VADU) SNPs. Importantly, Principal Component Analysis with the overlapping 472 SNPs or the 110 SNPs with identical profiles did not provide such clear demarcation (Supplementary Fig. 3, Additional File 2).

### Functional analysis revealed distinct biological processes in *Prakriti* groups across the cohorts

Next we asked whether there were any shared patterns of functional enrichments that differentiate the *Prakriti* types. Functional analysis using Gene Ontologies (GO) with the genes harboring differentiating SNPs in the three *Prakriti* groups comparisons revealed significant enrichments (p-value<10-2 without correction) (Fig. 2C) of biological processes in both the cohorts. For instance in (a) P vs K comparison we observed significant enrichment for specific ontologies such as Type I interferon and interferon-gamma mediated signaling pathways, cell movement or subcellular compartment and biological adhesion; (b) in the case of V vs K comparison significant enrichments for GO categories related to the regulation of synapse activity, regulation of cell development and anatomical structure formation, and (c) lastly, comparison of V vsP revealed specific enrichment for the processes like neurogenesis, positive regulation of nucleocytoplasmic transport, processes related to cAMP and carbohydrate derivative biosynthesis. These results are largely in line with the previous reports utilizing the gene expression profiles of NI Indian cohort [28,35].

**Fig. 1:**
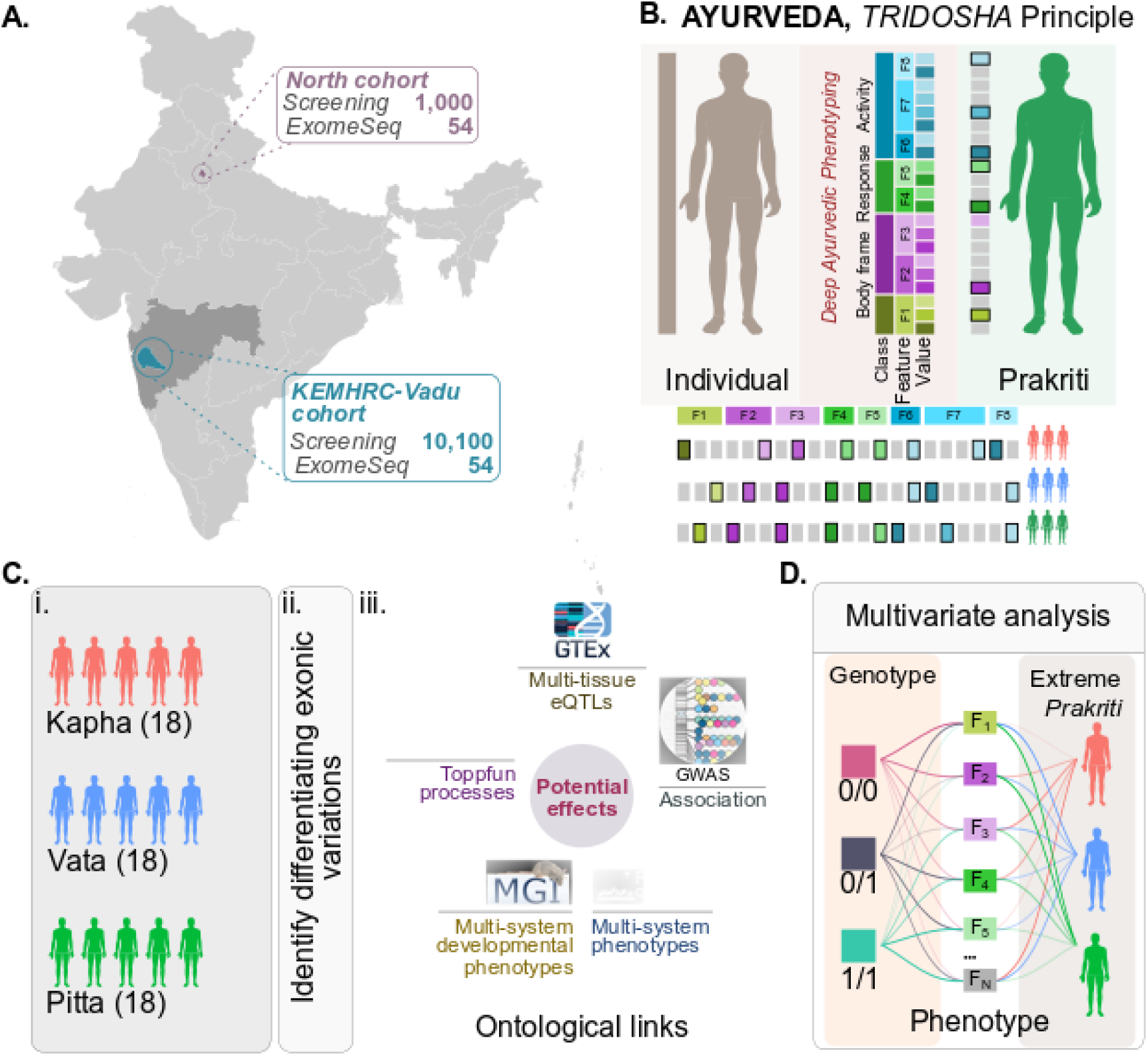
Schematic of the overall study. (a) Extreme *Prakriti* individuals were identified from two cohorts that belong to Indo-European linguistic lineage from Delhi (NI) and Pune, Maharashtra (VADU). These populations are members of a genetic cluster of Indo-European populations in the Indian Genome Variation Consortium (IGV). (b) Stratification of healthy individuals within a population using deep Ayurvedic phenotyping. *Prakriti* classification is based on a comprehensive assessment of ∼150 attributes that include anatomical, physiological, physical and psychological features. The three contrasting Vata, Pitta and Kapha *Prakriti* groups exhibit different phenomic architecture. (c) Workflow of exome sequencing and analysis is depicted. (i) The numbers of healthy individuals of each *Prakriti* groups from each cohort along with background populations included in the exome study is given in parenthesis. (ii) & (iii) Functional annotations of differentiating variants were carried using; Toppfun for Biological process annotation and enrichment analysis; GWAS for disease associations, GTEx for identifying cis-eQTLs, anchored expression of allelic states and effects on different tissues; PheWAS catalog for association with multisystem phenotypes and MGI mouse knockdown/knockout resource (MGI) for links to multisystem developmental phenotypes. (d) Multivariate analysis using random forest was used for inferring links from genotypes to *Prakriti* phenotype through the feature space using genotypes of replicated SNPs.

**Fig. 2:**
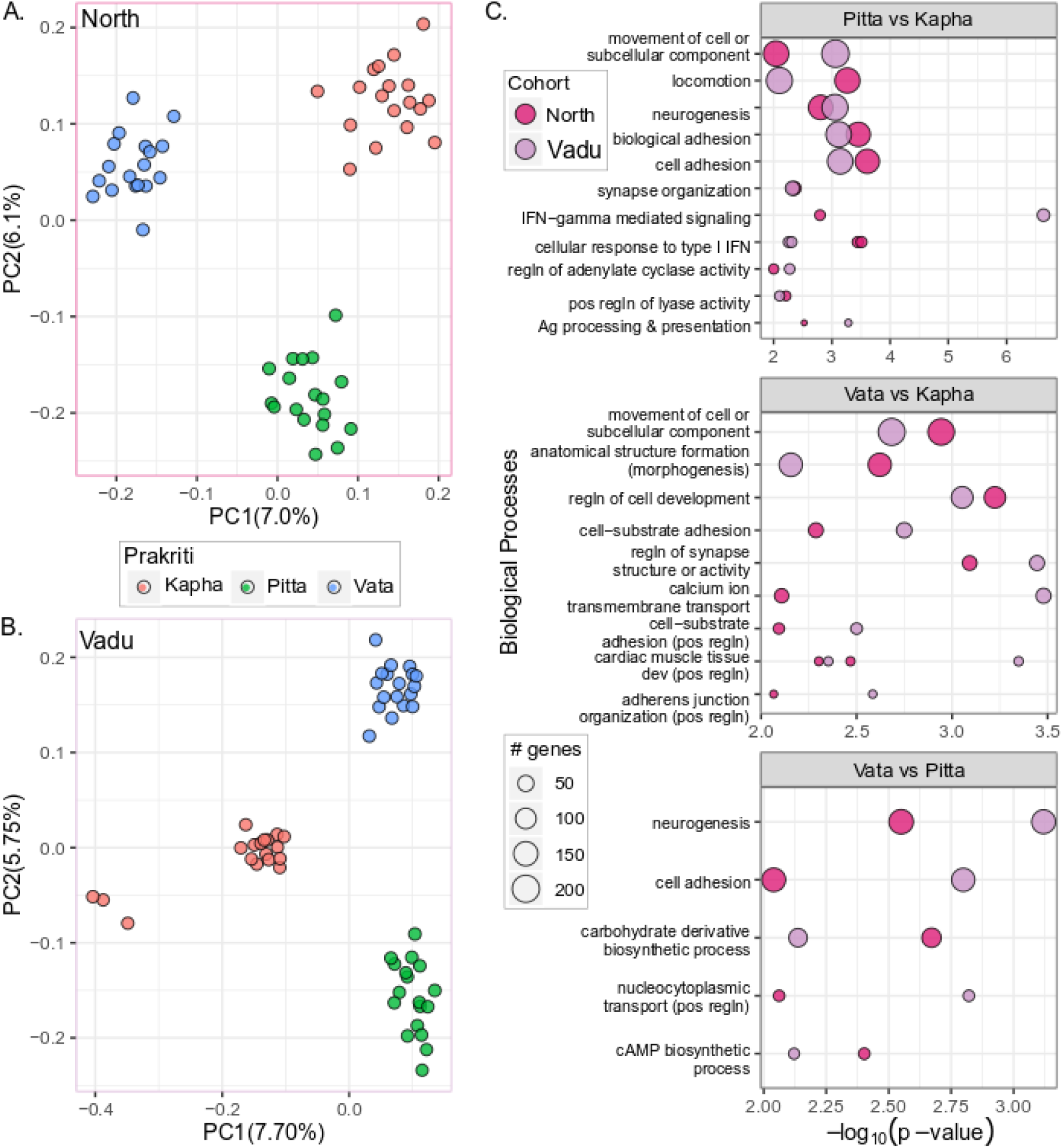
Differentiation of *Prakriti* groups based on SNPs and biological processes in both cohorts. (a) Principal Component Analysis (PCA) plot depicts segregation of *Prakriti* groups into three distinct clusters on the basis of SNPs from replicated set of 1181 genes in (i) NI and (ii) Vadu cohort. (b) Bubble plot depicts enriched (p-value<0.001) biological processes that are shared between the cohorts in each pair-wise *Prakriti* comparison P *vs* K, V *vs* K and V *vs* P; size of the bubbles indicate the number of genes in each process.

### *Prakriti* associated variants in common and complex diseases

Next, we investigated whether the differentiating SNPs are reported to be associated with complex traits in the GWAS catalog. We observe 119 and 166 SNPs that differ significantly between healthy individuals in the NI and VADU cohorts in the GWAS catalog (Supplementary Table 6, Additional File 1). Among these, six GWAS SNPs; rs8100241 (*ANKLE1*), rs2228145 (*IL6R*), rs2645294 (*WARS2*), rs1129555 (*GPAM*), rs2236293 (*TMEM8B*) and rs11084300 (*NLRP12*) were replicated in both the cohorts (Supplementary Table 6, Additional File 1). Besides, exact GWAS SNPs, we also observe ~11% of the differentiating SNPs in NI and VADU cohorts to be in strong LD (r 2>0.8) with GWAS SNPs. In many cases, different SNPs from the cohorts would tag the same GWAS SNPs or vice versa (Supplementary Table 7, Additional File 1).

The majority of these GWAS SNPs are also cis-eQTLs in GTEx data which allowed us to anchor the associated variants with expression across tissues and different *Prakriti* types. The differentiating variants between the *Prakriti* types that are reported in GWAS are also associated with multiple anthropometric traits of obesity, waist-to-hip Ratio (WHR), BMI and biochemical parameters that are measured in routine diagnostics for common diseases. This includes variants associated with BMI and WHR ratio adjusted for smoking behavior that differentiates Vata from Kapha; obesity-related traits and bone mineral density (TB-LM or TBLH-BMD) in Pitta and Kapha comparisons; and hand-grip strength, nose size, and hair morphology that differ between Vata and Pitta in both cohorts (Supplementary Table 6, Additional File 1). Variants associated with hematocrit parameters that are informative for health conditions related to anemia, infection and macrophage migration inhibitory factor levels are mostly seen in Vata comparisons (Supplementary Table 6, Additional File 1). SNPs were observed in genes associated with traits that are distinguishing features of the *Prakriti* types such as skin pigmentation, circadian rhythm, sleep functions, and sensory perceptions [27]. We also observe variants associated with disease susceptibilities which include allergy and infection prominently in VvsK and VvsP comparisons, inflammation in PvsK and VvsP and neuropsychiatric conditions in VvsK. Variants associated with a cerebrospinal fluid biomarker (rs2228145, *IL6R*)[36] differentiate Vata and Kapha in both cohorts. rs2228145 polymorphism in *IL6R* also influences the function of IL6-a pro and anti-inflammatory cytokine. Alternate allele -C-carriers of this SNP have decreased inflammatory response as well as the decreased prevalence of metabolic syndrome, diabetes, and atrial fibrillation.

Amongst the replicated SNPs, *NLRP12* variants share the same profiles in *Prakriti* comparisons. *NLRP12* gene regulates the immune system’s response to injury, toxins or invasion by microorganisms. This gene, unlike most NLR proteins, inhibits the release of certain molecules during inflammation. The replicated variant rs11084300 in *NLRP12* is associated with macrophage migration inhibitory factor levels [37] and differentiates Vata from Pitta. Taken together this suggests that integration of *Prakriti* based methods in healthy individuals could complement genomics-based risk stratification for complex diseases for early interventions.

### *Prakriti* differentiating variants in PheWAS studies

*Prakriti* differentiating genes and SNPs that map to PheWAS catalog suggest plausible pleiotropic effects. We observe nine SNPs in NI and 12 in the VADU cohort that associate with multi-system phenotypes in the PheWAS catalog (Table 1). Also, 83 and 66 *Prakriti* differentiating SNPs are in strong LD with 45 and 43 PheWAS SNPs in NI and VADU cohort respectively (Supplementary Table 8, Additional File 1). The replicated GWAS SNP rs8100241 in *ANKLE1* that is associated with ovarian & breast cancer is in strong LD with rs2363956 from the PheWAS Catalog. This SNP associate with multiple phenotypes in the PheWAS catalog (Supplementary Fig. 4, Additional File 2). Five replicated SNPs from *ATP5SL* locus associated with height in the GWAS catalog are in strong LD with the PheWAS SNP rs17318596. The associated multi-system phenotypes include intestinal infection, viral infection, obesity, and other disorders of metabolic, endocrine and immune systems.

**Table 1:**
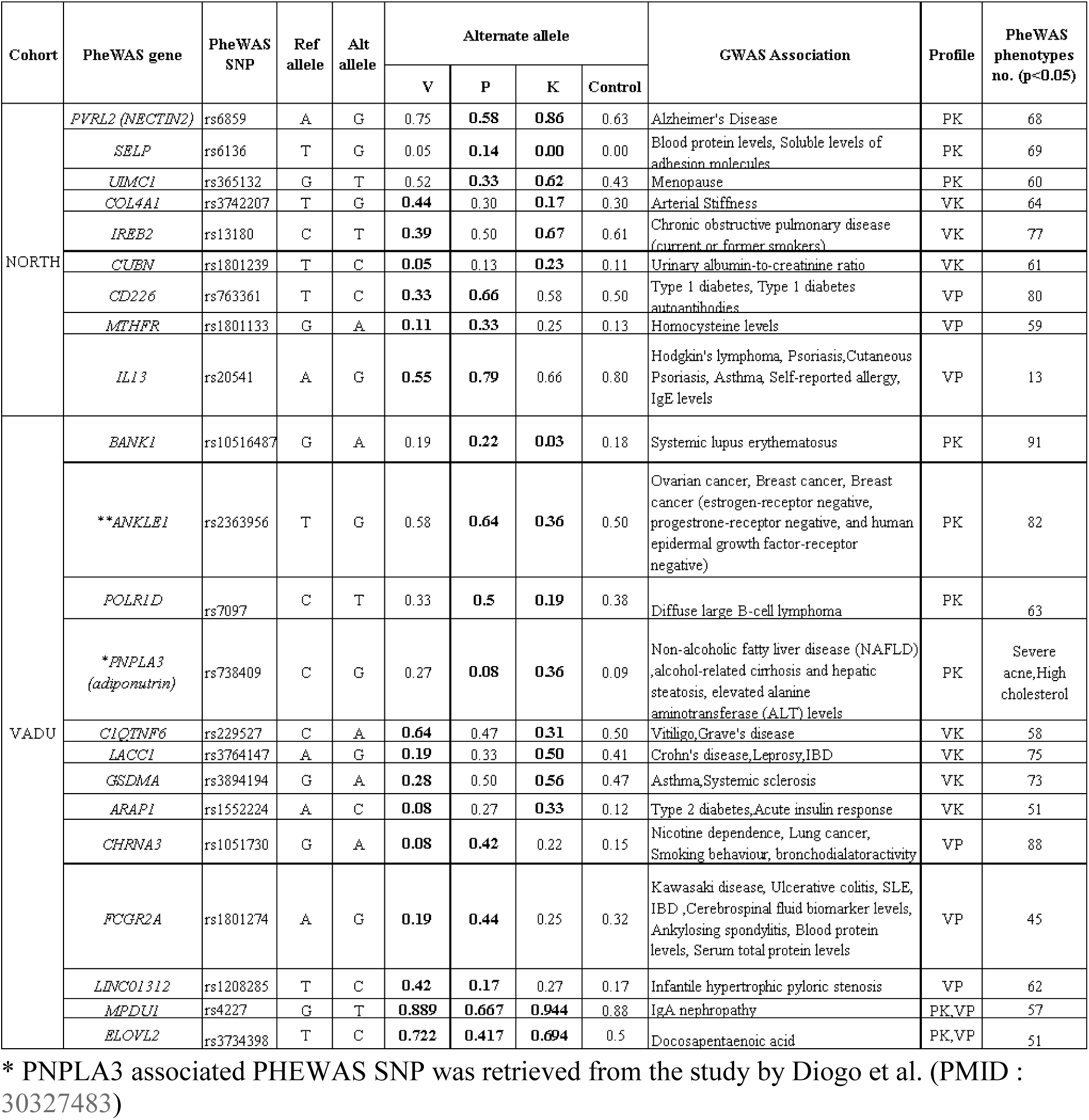
*Prakriti* differentiating SNPs reported in PheWAS catalogues in both cohorts.

### Potential pleiotropic outcomes of differentiating variants on gene expression and developmental phenotypes

The Mouse Genome Informatics (MGI) database has earlier been used to study the multisystem effect (physiology and phenotype) of mutations in orthologous genes reported in GWAS [38]. We infer the potential consequence of the genotypic states of the variants through mining information vis-a-vis association with expression across diverse tissues in GTEx [39] and/or phenotypic consequence in orthologous gene knockdown/knockout in mouse [40]. More than 80% of differentiating variations from both cohorts are eQTLs in GTEx data (Supplementary Table 10, Additional File 1) of which 30% to 70% map to multiple tissues, suggesting their potential effect at the system-wide level (Supplementary Fig. 5, Additional File 2). The differentiating SNPs are most enriched in nerve tibial, skin sun-exposed lower leg, testis, thyroid and subcutaneous adipose tissue. A substantial fraction of the SNPs also affects adjacent genes. The proportion of eQTLs observed in each *Prakriti* comparison group is correlated (r^2^>0.98) between the cohorts.

91 out of 110 replicated SNPs are eQTLs with 39 having effect size less than −0.4 or greater than 0.4 and p-value<10^−7^(Supplementary Table 9, Additional File 1). We demonstrate examples of two replicated SNPs from *IFIT5* and *ZNF502* genes that map to anti-viral response (Supplementary Table 3, Additional File 1). *IFIT5*, a member of IFN-induced protein with tetratricopeptide repeats which enhances innate immune response during RNA virus infection differs significantly between Pitta and Kapha. Both the SNPs are eQTLs in GTEx with prominent effect sizes in diverse tissues like salivary gland, spleen and colonic tissue (Fig. 3A; Supplementary Fig. 6A, Additional File 2). The alternate allele -C-of rs304447 that associates with lower expression are significantly depleted in Pitta compared to Kapha. A similar pattern with respect to the immune response is observed in another replicated SNP rs56084453 that maps to *ZNF502*. The alternate allele of rs56084453 that associates with significant downregulation of *ZNF502* have a prominent effect size in the spleen, salivary gland and ileum (Fig. 3B); Supplementary Fig. 6B, Additional File 2). This allelic state is fixed in Pitta in both cohorts and differs from Kapha significantly. Both these observations suggest that variations associated with Pitta could be involved in enhanced anti-viral response.

**Fig. 3:**
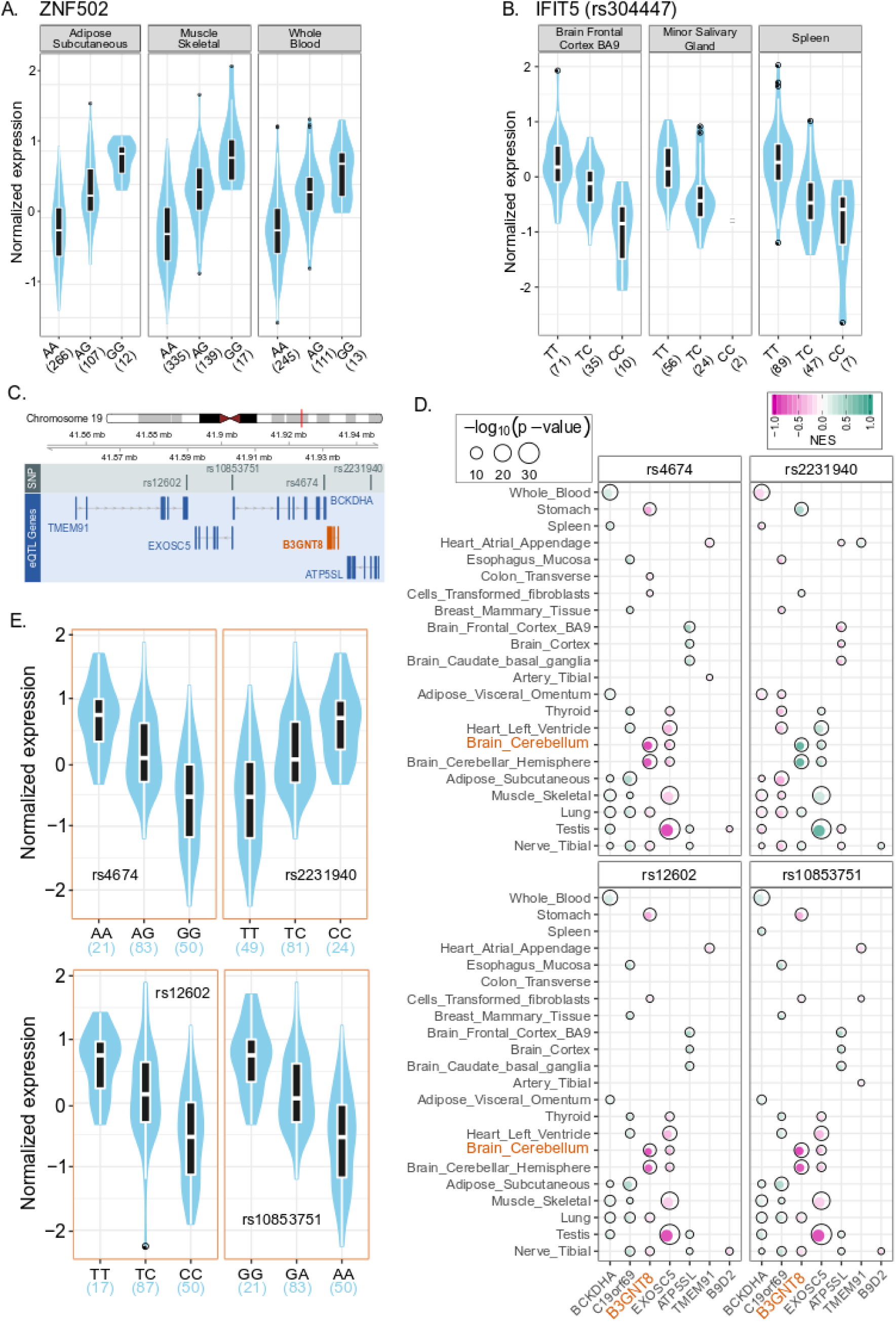
Tissue-wise effect of replicated SNPs (with same profiles in both cohorts) on gene expression in GTEx v7 data. Violin plots of normalized expression across representative tissues are depicted. Effect size and bubble plots of expression across all GTEX tissues are provided in (Table S9 and Fig. S6). (a) Violin plots of rs56084453 genotypes of *ZNF502*. The alternate A allele is associated with downregulated transcripts. (b) Violin plot of rs304447 in *IFIT5*. The alternate allele C of rs304447 in IFIT5 is associated with its lower expression. Frequency of alternate allele is significantly lower in Pitta group than Kapha. (c) (i) Schematic representation of a 54kb region in chromosome 19 harbouring multiple Vata and Kapha differentiating SNPs rs12602 (*TMEM91*), rs10853751 (*EXOSC5*), rs4674 (*BCKDHA*), rs2231940 (*ATP5SL*) in GTEx tissue that are replicated in both cohorts. (ii) Bubble heatmap depicts the significance and effect size of each of the eQTLs on multiple genes in the same locus. All the eQTLs have maximum effect on *B3GNT8* across different tissues. Elevated levels of *B3GNT8* have been shown to be associated with glioma. (iii) Violin plot depicts the effect of all the above SNPs on *B3GNT8* expression in cerebellum.

We had an important observation wherein region spanning 54 kbps with four replicated SNPs in overlapping genes; *BCKDHA, TMEM91, EXOSC5* and *ATP5SL* that differ in a *Prakriti* specific manner. The frequency of the alternate alleles of rs4674 (*BCKDHA*), rs12602 (*TMEM91*) and rs10853751 (*EXOSC5*) and reference allele of rs2231940 (*ATP5SL*) are lower in Vata compared to Kapha in both the cohorts (Supplementary Table 3, Additional File 1). All these SNPs are eQTLs and exert effects on each other as well as adjacent genes across a large number of tissues (Fig. 3C-E)). Noteworthy, the Vata associated allelic state has a significant positive effect (p<10^−10^ with effect size 0.89) on an interspersed gene (*B3GNT8*) in the brain cerebellum (Supplementary Table 9, Additional File 1).

Out of the 87 genes with replicated SNPs and similar patterns of difference between the two cohorts, mouse knockdown/knockout phenotypes exist for 40 genes (Supplementary Table 12, Additional File 1). We used information on mouse phenotypes coupled with functional annotations of these genes in Toppfun to infer their involvement in biological and cellular processes and potential to affect different phenotypes in mouse orthologs. As seen in alluvial plots for the 40 genes (Fig. 4A), most of these genes could potentially impact processes at different functional hierarchies as well as multiple human phenotypes that are captured during *Prakriti* assessments. We further illustrate in detail the *EDNRA* gene, which encodes the receptor for endothelin-1, a peptide that plays a role in potent and long-lasting vasoconstriction. Variability in this gene is likely to impact a large number of cellular and physiological processes and affect functioning of different organs and systems. These could manifest as syndromic features in healthy individuals that differentiate the *Prakriti* types (Fig. 4B). Similar to *EDNRA*, the other genes with large effect, *SIX1*, *CHD5*, *FIG4* and *HNRNPD* are involved in number of cellular and biological processes and affect multisystem phenotypes (Fig. 4A). These include many anatomical features such as body frame, body build, skin texture, physiological attributes like metabolic patterns, voluntary and involuntary movements (Fig. 4B; Supplementary Fig. 7, Additional File 2).

**Fig. 4:**
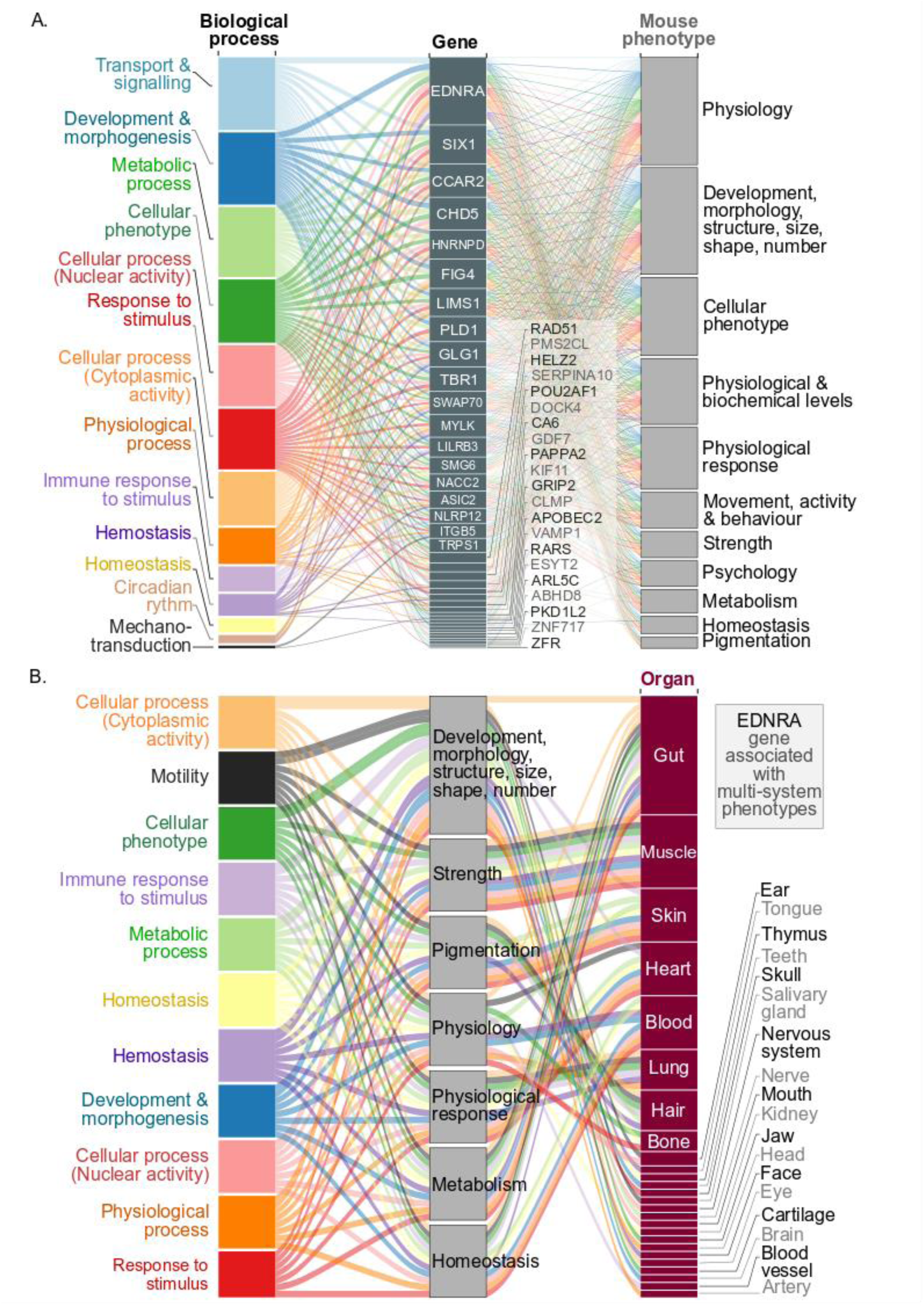
Alluvial plot representing potential impact of *Prakriti* associated genes on biological processes and mouse developmental phenotypes. (a) Alluvial map of 40 genes that have replicated SNPs with identical profiles in both cohorts. (b) Alluvial plot of *EDNRA* gene and its connectivity to various biological processes and mouse phenotypes suggests inherent variability in this key vascular endothelial receptor gene could have a multisystem impact.

### Extreme *Prakriti* types could identify variants with pleiotropic effects

Next, we wanted to infer whether Prakriti associated GWAS variants could have pleiotropic effects like in PHEWAS studies. We inferred this through analysis of variant effects on expression in GTEx [39], phenotypes in mouse knockout/knockdown [40] and PheWAS catalog [12]. SNPs in *ANKLE1* (rs8100241 & rs8108174) and *ABHD8* that differ between Vata and Kapha are in LD with each other and maps to a modifier pleiotropic locus implicated in estrogen negative breast cancer in *BRCA1* carriers and progression in ovarian cancer [41]. *ANKLE1*variations have been shown to have a negative effect on *ABHD8* expression which in turn enhances invasiveness [41]. In GTEx the differentiating SNPs in *ANKLE1* is observed to lower the expression of *ABHD8* significantly in breast mammary tissue, tibial artery and nerve, esophagus mucosa, skeletal muscle, subcutaneous adipose, skin in sun exposed lower leg and testis. The allelic variant in *ANKLE1* linked to lower *ABHD8* expression is over-represented in Vata. This suggests a more invasive molecular phenotype could be associated with Vata *Prakriti*. Further, *ABHD8* knockdown phenotypes in mouse exhibit features such as decreased body length, bone mineral content and density, short tibia, lean body mass, hyperactivity and increased exploration in new environment. These features resonate with Vata phenotypes. The replicated SNP in *ANKLE1* (rs8100241 & rs8108174) are also reported to be in LD with a PheWAS SNP (rs2363956) and besides neoplasms has been associated with phenotypic attributes such as mood disorders, loose joints, involuntary movements and Atrioventricular (AV) block (Supplementary Fig. 4, Additional File 2).

A recent study conducted in four large cohorts with extensive health records (700,000) from 23 & Me, UK Biobank, FINRISK, CHOP identified novel phenotypic association of 19 existing drug targets identified in GWAS[16]. A validated variant from this large scale study in *PNPLA3* (rs738409) was also observed to differ between *Prakriti* types in VADU cohort. The -G-allele of *PNPLA3* is a potential drug target for alcohol-related cirrhosis, Non-Alcoholic Fatty Liver Disease (NAFLD) and hepatic steatosis. GWAS studies also associate with severe acne, high cholesterol, anti-cholesterol medication, gout and gallstones. The -G-allele is significantly over-represented in Kapha and -C-allele in Pitta *Prakriti*. The traits that have been associated with the allelic states resonate with the phenotypic and susceptibility differences between Pitta and Kapha. Noteworthy, amongst them is severe acne that is a distinguishing feature of Pitta *Prakriti* [28,35]. Recently, WES of NAFLD patients with extreme phenotypes reconfirmed the involvement of this variation in progression to fibrosis [42].

Another striking example was of the SNP rs2228145 in *IL6R* gene. Clinical trials of IL6 receptor blockade using antibodies, tocilizumab and sarilumab, have revealed aortic aneurysm and atopic dermatitis as notable side-effects [14]. Functional studies of this natural variant of *IL6R* receptor have revealed that carriers (-A-allele, Asp328Ala) have lower expression of *IL6R* [43,44]. The - A-allele of the *IL6R* variant has been associated with the risk of rheumatoid arthritis and the -C-allele with atopic dermatitis and allergic diseases in GWAS. A PheWAS study conducted on a million veterans program and replicated in UK Biobank and Vanderbilt biobank (800K) have reported new associations of aortic aneurysms as well as arthritis with the -A-allele and atopic dermatitis with the -C-allele [14]. We find that the -A-allele differentiates Vata from Pitta and Kapha in NI and VADU cohort respectively. Analysis in these genes, *ANKLE1*, *ABHD8*, *PNPLA3*, and *IL6R* suggests that *Prakriti* stratification could help segregate individuals with different disease outcomes and susceptibility. Thus, holds potential to identify individuals with minimal risk associated with *IL6R* antibody treatment for several chronic inflammatory conditions.

### Genotypes to composite *Prakriti* types through phenotype feature space

As *Prakriti* exhibits phenotype-phenotype connectivity, we investigated the plausible linkage of the differentiating SNPs with minimal features; and the power of minimal features to predict *Prakriti* types. We carried out a multivariate analysis using random forest [45] to infer potential interactions. This experiment allows us to decipher the likely dependency of SNPs through features on *Prakriti* types. With an error class rate set at 30% and despite our limited cohort set, we could identify important interactions with 84 (7.7%) and 99 (6.8%) SNPs in NI and VADU cohort, respectively, have linkage from genotype to *Prakriti* through features space (Supplementary Table 10, Additional File 1). Most SNPs 965 (88.6%; NI) and 1256 (86.7%; VADU) were not found to have direct association with features that can predict *Prakriti* types. Interestingly, 40 (3.7%) and 94 (6.5%) SNPs could be predicted using some features, but these features were inadequate in predicting *Prakriti* types. The complete list of SNPs with their predictive power with features (G2F) and corresponding features to *Prakriti* types (F2P) is provided (Supplementary Table 10, Additional File 1).

One noteworthy observation was SNP rs7244213 in the urea transporter gene *SLC14A2*. The homozygous reference allele, and the heterozygous clearly distinguished attributes for 17 features related to anatomical, physiological and physical activities (Fig. 5; Supplementary Fig. 8, Additional File 2). This ensemble of features also differentiated Kapha from Pitta and Vata in the heterozygous state. These sets of features are the most distinguishing features of Kapha from Pitta and Vata groups as described in Ayurveda text (Supplementary Information).

**Fig. 5:**
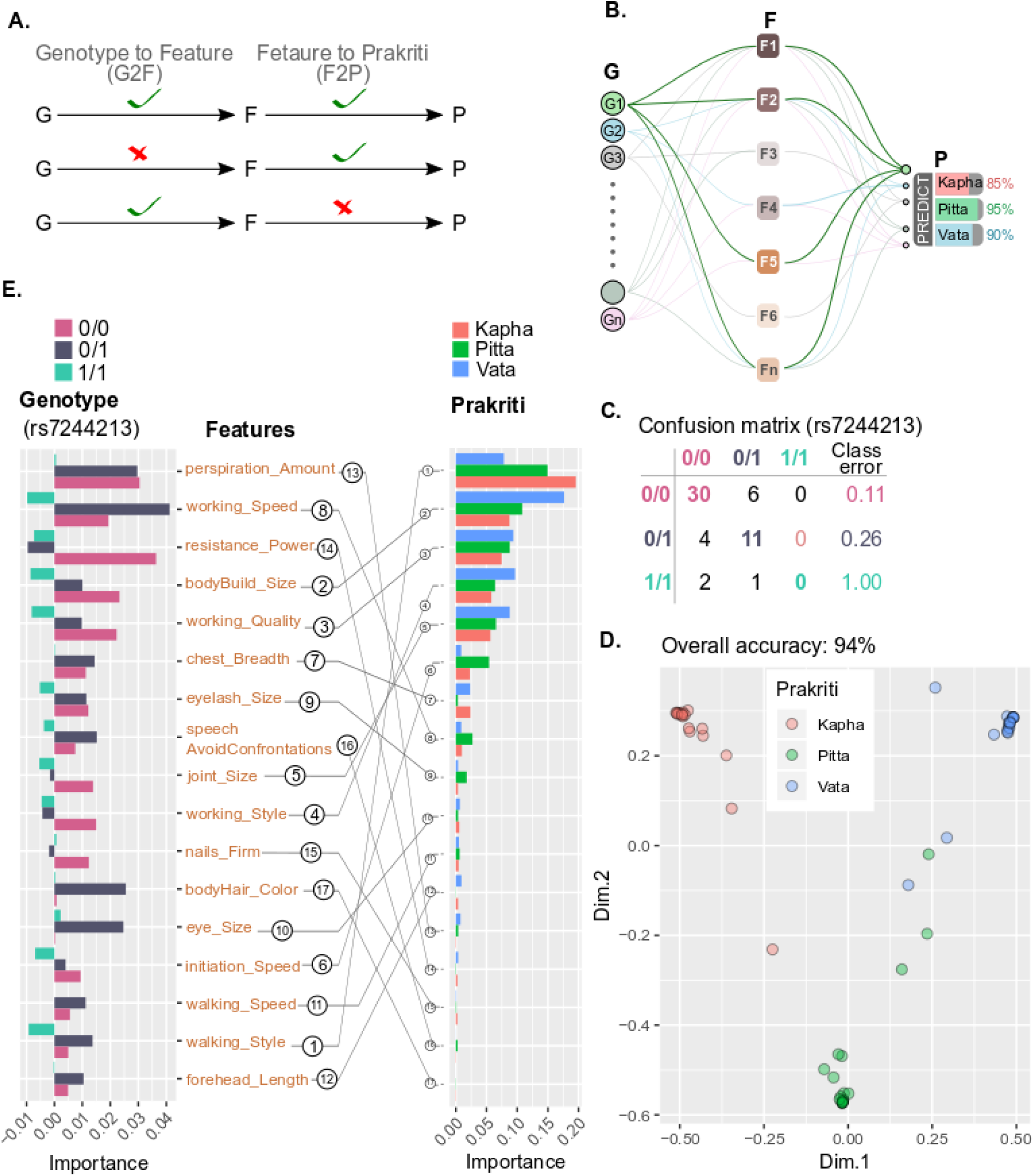
Random forest (RF) analysis to probe the connectivity of genotype to *Prakriti* through features (deep phenotypes) captured in questionnaire. (a) Schematic of possible associations of genotype links to (i) features, (ii) *Prakriti* through features space or (iii) *Prakriti* that can be derived from existing data. (b) Schematic of the RF analysis. This was carried out for each genotypes (replicated SNPs) using∼150 features captured during *Prakriti* assessment. Based on the model performance of 70% overall accuracy, we selected our best models. (c) (i) Confusion matrix for the genotypes of rs7244213 based on RF model with17 best features. This SNP maps to a urea transporter *SLC14A2* involved in osmoregulation. Feature importance plot of 17 predictors of (ii) rs7244213 (*SLC14A2*) and (iii) *Prakriti*. Order of features is based on importance from the genotype model and serial number depicts the rank of these features in the *Prakriti*. (iv) Multi-Dimensional Scaling (MDS) plot derived from RF model using 17 predictors of rs7244213, to classify *Prakriti*.

## Discussion

It is anticipated that identification of genetic variants with pleiotropic effects that can contribute to heterogeneity in health and disease outcomes will strengthen personalized medicine. Phenome-wide association studies (PheWAS), which analyze multiple phenotypes in relation to the genotype from genome-wide association studies have been successful in identifying such genetic variants. Here we outline a novel alternative to PheWAS approach: the identification of pleiotropic variants through analysis of multi-system composite phenotypes classified by Ayurveda as extreme constitution types, through exome sequencing.

We wanted to explore whether this approach could help identify common variations that determine health and disease outcomes and link to composite phenotypes. Strength of our study includes use of in-depth composite and extreme phenotypes and small sample sizes. Since Ayurveda describes *Prakriti* on the basis of underlying biological variability, we investigated whether the genetic differences provide meaningful insights. We therefore studied the overall patterns of differences with respect to functional outcomes and links to biological descriptions of *Prakriti*. A core set of differentiating SNPs from 1181 replicated genes could clearly distinguish the three groups in both the cohorts with 110 SNPs from 87 replicated genes having similar profiles in *Prakriti* comparisons. These genes govern different biological processes and are associated with multisystem attributes in mouse knockdowns/knockouts, few noteworthy ones being *EDNRA*, *SIX1*, *FIG4* and *KIF11*. Many of these features overlap with phenotypic attributes that differentiate *Prakriti* types (Supplementary note; Supplementary Fig. 9, Additional File 2) that suggest inherent variability in these genes could confer phenotypic differences amongst *Prakriti* types.

It is noteworthy that the three distinct *Prakriti* types cluster on the basis of 1181 replicated genes despite limited overlap in the SNPs. The *Prakriti* differentiating genes map to distinct biological processes of immune response, cell adhesion, motility, anatomical structure development, signaling and chemosensory perceptions across cohorts. This corroborates with our earlier observations on genome-wide expression differences amongst the *Prakriti* groups [28]. It also highlights the utility of such a phenotyping approach for sub-stratification in genetically diverse populations. When we anchor the likely effects of allelic variants on expression in GTEx we find 80% of them to have substantial effects in multiple tissues. The patterns of tissue-specific enrichment of eQTLs are also similar between the cohorts. Taking a few examples of genes from enriched pathways we show how these differences could be meaningful for *Prakriti* specific outcomes. For instance, eQTL in genes from innate immune response pathways such as *IFIT5*, *ZNF502* suggests Pitta *Prakriti* individual might counter viral infections more efficiently [28]. In a recent study it has been shown that knockdown of *ZNF502*, limits replication of the human respiratory syncytial virus (RSV) [46]. The natural variation rs56084453 associated with lower expression of *ZNF502* that is fixed in Pitta in both cohorts might, therefore, confer protection from recurrent viral infections (Supplementary Table 10, Additional File 1).

Another noteworthy observation was a 54 Kbps region encompassing the replicated SNPs covering four genes, *BCKDHA*, *TMEM91*, *EXOSC5* and *ATP5SL* that appears to have evolved for a functional requirement. This is retained as a haplotype in *Prakriti* groups and directions of expression of all the variants are also constitution specific (Fig. 3C, Supplementary Table 11, Additional File 1). All the eQTL variants from Vata are associated with higher expression of *B3GNT8* in the brain cerebellum, elevated levels of which govern the metastatic potential of many cancers most prominently glioma [47].

We make similar observations with respect to the other modifier locus harboring *ANKLE1* and *ABDH8* that is reported to be a pleiotropic locus governing variability in the progression of breast and ovarian cancer [41]. These observations suggest that sub-stratification in disease conditions using *Prakriti* methods might help identify endophenotypes for stratified interventions.

In our exome analysis we also observed a large number of SNPs amongst healthy individuals stratified into extreme *Prakriti* groups that had exact matches or were in LD with the GWAS and PheWAS associated variants. Our results suggests that estimation of genomic risk scores in Biobanks and prospective cohorts if carried out in conjunction with phenotypic stratification of *Prakriti* could enable identification of endo-phenotypes with different health and wellness management needs. Conditioning genetic association with *Prakriti* information in rheumatoid arthritis and pharmacogenetics have earlier highlighted the merit of such an approach [48].

Inferences of systemic effects of *Prakriti* differentiating variations from reported literature prompted us to further explore the interaction of genotypes using the wealth of phenotypic attributes and its connectivity that characterizes the *Prakriti* type. Random forest analysis using SNPs from replicated genes with the individual’s phenotypic features revealed cryptic genotype to phenotype links. An interesting observation was the variant in the urea transporter *SLC14A2* which is an important component of the hypothalamic-neurohypophyseal-renal axis that is involved in the maintenance of water balance during variations in water intake as well as in-utero during fetal development [49]. Amongst the osmoregulation related genes, *SLC14A2* has been shown to be under adaptive evolution in cetaceans during evolutionary transition from a terrestrial to a hyperosmotic environment [50]. Further, the differentiating variation rs7244213 has also been associated with hypertension in East Asians [51]. Genotypic states of this SNP in *SLC14A2* associate with an ensemble of 17 features that distinguishes the three *Prakriti* types (Fig. 5; Supplementary Fig. 8, Additional File 2). Some of these phenotypes can also be attributed to the water balance function of this gene inclusive of amount of perspiration and body frame. This observation is important for biological understanding of Vata and Kapha functions reflected in *Prakriti* features governed by their dry and humid nature respectively. There are specific therapeutic strategies described in Ayurveda for restoring Vata-Kapha balance through the management of fluid balance [28,29]. This genotype-phenotypic association though extremely preliminary, provides a novel window to explore water balance disorders. Small-molecule inhibitors against these urea channels are being proposed as a new class of aquaretics with potential for their usefulness in hyponatremic disorders [52]. Our observations suggest that such inhibitors might have variable requirements and outcomes amongst individuals of different *Prakriti* types.

A limitation of the study may be the low sample size. However since this is a pilot study, we also estimated the power of study based on the allele frequencies as well as sample requirements for future adequately powered study designs (Supplementary Table 4, Additional File 1; Supplementary Fig. 2A-B, Additional File 2). Using different p-value cutoffs, simulation studies based on the frequency differences of the differentiating SNPs, we estimate substantial power if the studies are conducted even in 50 samples of each group (alpha=0.05) in both the cohorts. Given the composite nature and phenotypic architecture of *Prakriti*, we might need to evolve new methods for estimation of power and our observations though preliminary, would be useful for such calculations. Noteworthy, power estimation methods are still not the state-of-the-art for PheWAS [12].

## Conclusion

Sequencing of healthy individuals of extreme phenotypes of *Prakriti* has provided (a) a set of genes that can cluster *Prakriti* groups and have the potential to predict differences in disease progression, response to environmental triggers and therapeutic interventions (b) composite traits in constitution specific manner that could enable deep-phenotyping for stratification of healthy and diseased individuals. Further the genotype-phenotype connectivity in a *Prakriti* specific manner can be formalized through machine learning approaches. We anticipate that integration of this framework in existing case-control studies, Biobanks and prospective cohorts could increase the yield of genes with pleiotropic effects and provide an ensemble of features that enable identification of target populations for precision interventions.

## Materials and methods

### Sample description

Exome study was carried out on healthy subjects of predominant *Prakriti* types identified from our two earlier studied cohorts. This include 108 (18 x 3 x 2) individuals with 18 each Vata (V), Pitta (P) and Kapha (K) in each cohort. An extensive protocol was followed for recruitment of subjects, clinical phenotyping, classification into predominant groups as well as establishment of genetic homogeneity as previously published [28,29].

These *Prakriti* types comprise 10% of the studied population, belong to the age group of 18-40 years, exhibit differences with respect to 150 multisystem features that include anatomical and physical attributes as well as physiological and psychological responses [35]. Genetic homogeneity of the study cohorts and its relatedness to diverse Indian population was affirmed by principal component analysis using a set of 17675 SNPs that overlaps with the Indian Genome Variation Consortium (IGVC) diversity panel [53]. The study has been carried out as per protocols approved by institutional human ethics committee at CSIR-Institute of Genomics and Integrative Biology, Delhi and KEM Hospital Research Centre, Pune, India.

### Whole Exome Sequencing (WES) and Variant Calling

Exome sequencing of 108 healthy subjects was carried out on Illumina HiSeq2000 platform using standard methods. GATK Best Practices Workflow was followed for processing sequencing reads and calling variations. Variants with less than 50% genotyping call rates or quality scores less than 30 were removed prior to analysis. We used Vcftools 0.1.12 to convert genotypes in VCF format to Plink format for statistical analysis. Variants were annotated using Annovar [54] with novel variants indicated by their chromosomal position in version GRCh37. We carried out three pair-wise comparisons; Vata *vs* Pitta (V *vs* P), Pitta *vs* Kapha (P *vs* K) and Vata *vs* Kapha (V *vs* K) to identify differentiating variants. We used Fisher’s exact test (p-value<0.05) implemented in PLINK (v 1.7). To assess whether the differences were *Prakriti* specific, we carried out permutation analysis by randomly shuffling the *Prakriti* labels ~50,000 and ~80,000 times per SNP for NI and Vadu cohort respectively. The numbers of iterations were based on the number of significant SNPs in pair-wise comparison at FDR of 5%. The SNPs that were present in lower 5% distribution of p-values of the permuted set were retained. Profiles are indicated on the basis of alternate allele frequencies, for example, V-K+ indicate the lower frequency in Vata in V *vs* K comparison.

As the study involves extreme and composite phenotypes that comprise 10% of the population, we anticipated adequate power in smaller sample sizes. However, no estimates of sample sizes, for an adequately powered study on extreme and composite types, that too involving only healthy individuals are available. We therefore estimated the power of the study based on the allele frequencies observed from our data on two cohorts from each comparison group (e.g.,Vata *vs* Kapha). We quantified power using a power.fisher.test function of statmod [55] R-package. Power estimations were done using simulations performed with increasing sample sizes; original sample numbers used for frequency estimation, 18, 50, 100, 500, 1000, 10,000, 50,000 and100,000 with alpha of 0.05. Additionally, we estimated power with constant sample size (N=500) and varying alpha ranging from 10^−2^ to 10^−12^ for each SNP with their corresponding allele frequencies.

### Replication Analysis of *Prakriti* differentiating SNPs

The extent of replication was assessed at three different levels of genes (1) with identical and/or different SNPs as well as profiles (2) with identical SNPs having similar and/or different profiles (3) identical SNPs with exactly matching profiles in both cohorts. Principal component analysis (PCA) using EIGENSTRAT [56] was carried out to assess the extent of differentiation between the *Prakriti* on the basis of significant SNPs from only the replicated genes. Top 20 principal components were identified for variance estimation.

### Functional Annotation of differentiating variants

Gene Ontology (GO) annotation of differentiating SNPs from each *Prakriti* group comparison was carried out using Toppfun[57]. The allele-specific consequences were assessed using the Genotype-Tissue Expression (GTEx) [39] Project v7 data. We queried for tissue-specific cis-eQTLs (p-value<10^−7^& effect size < or >0.4). We also queried the SNPs for disease associations using (https://www.ebi.ac.uk/gwas/) GWAS catalog v1.0.2 (associations\_e93\_r2018-08-14). We used liftover to convert the exome coordinates (GRCh37) to GRCh38 coordinates. The South Asian (SAS) population in the 1000 genomes database was used for mapping GWAS SNPs in LD (r^2>0.8) with our variants using SNIPA [58]. The GWAS exact as well as proxy SNPs from the exome data were used to query PheWAS (p-value<0.05, uncorrected) catalog. Literature resources were also used for identifying variants with pleiotropic effects.

To explore the links between variability in replicated gene with functions and multisystem phenotypes, we curated the information from ToppGene Suite [57] and The Mouse Genome Database (MGD; http://www.informatics.jax.org) [40]. We categorized the processes into major groups of development & morphogenesis, physiological process, metabolic process, homeostasis, cellular process (nuclear activity and cytoplasmic activity), cellular phenotypes, transport and signaling, immune response, response to the stimulus, mechanotransduction, hemostasis, and circadian rhythm. The mouse phenotypes were grouped on the basis of anatomical, physiological, activity and behavior, and also the organs and tissues in which these were observed. Alluvial maps were generated using alluvial package [59].

### Multivariate Analysis for exploring genotype to *Prakriti* links through phenotype feature space

We carried out a multivariate analysis to assess whether the links to the differentiating SNPs to *Prakriti* are through the composite set of features that are used during assessment. We carried this analysis on 1181 replicated genes in both cohorts. After removing genotypes with missing values we used 1084 and 1449 SNPs from NI and VADU cohorts respectively. In the first step, we identified genotypes that could be explained through a composite set of minimal features (model genotype, predict genotype with features) with the least accuracy of 70% or more. These minimal set of features for a given SNP were used to predict the *Prakriti* types (model *Prakriti*, use minimal features to predict *Prakriti*) in the next step. The data were binned into following three plausible groups of SNPs which show:

i. Complete linkage from genotype to minimal features and minimal features to *Prakriti*.
ii. Partial linkage, where genotype could not be predicted by features, but feature to *Prakriti* linkage exist.
iii. Partial linkage, where genotype could be predicted by minimal features, but these features could not predict *Prakriti*.

We built classification models with random forest (RF) algorithm [45] to predict SNPs using the questionnaire and genotype data of both cohorts. The model building was performed independently for both cohorts. For a given SNP, genotype states (2 or 3) such as (A/A and T/T; or A/A, A/T and T/T) were used as class (Y) and questionnaire data were used as features (X). We performed feature selection using the Boruta algorithm with 500 iterations [60]. For each model, we used 20000 decision trees and the number of variables to be sampled at each node (mtry), was set as default (square root of the total number of variables). Based on the class-error rate less than 30% in minimum of two classes, we filtered the SNPs and extracted their corresponding best features from the RF model. To understand the power of these features for *Prakriti* prediction apart from the genotype we built random forest models without feature selection with 20000 decision trees.

## Acknowledgments

Authors acknowledge the staff and study population from both NI and KEMHRC-VADU, Pune, areas for their participation in the study. Project funding MLP-0901 from Council of Scientific & Industrial Research and CSIR-IGIB for administrative, infrastructure and IT support including the high performance computing facility. Authors acknowledge Raghu Nanadanan MV for IT infrastructure and data management of TRISUTRA data. Authors acknowledge Greg Gibson from Centre for Integrative Genomics, Georgia Tech University Atlanta for critical inputs and review and Saurabh Ghosh, Department of Genetics from Indian Statistical Institute, Kolkata is duly acknowledged. Authors acknowledge Ashish Kumar from Swiss Tropical and Public Health Institute, Basel, Switzerland for critical inputs and review and Megha from University of Transdisciplinary Health Sciences and Technology, Bangalore and Gaurav Ahuja from IIIT Delhi for help with editing of the manuscript. UGC Fellowship to TA, GC and TRISUTRA project fund support to RP, RK, GB and AN is duly acknowledged.

## Author’s contributions

M.M. and B.P. designed the project; R.P., G.C. performed NGS sequencing; T.A., P.D., V.M.,A.N., M.M., D.D., performed NGS analysis; T.A., R.K., A.N., R.P., P.S., G.B., D.D., B.P., MM performed data analyses; T.A., R.K., A.N., R.P., D.D., B.P., M.M. wrote the manuscript; R.K. and T.A. contributed to data visualization; R.K. performed machine learning; B.G., A.S., B.P., Ru.P., D.A., and S.J. phenotyping, sample collection and processing; M.M., D.D., R.P., and B.P. reviewed the data and manuscript. All authors have read and approved the manuscript.

## Additional Files 1

### Supplementary Tables

All the supplementary tables were uploaded at zenodo because of its large file size.

**1. Table S1:***Prakriti* differentiating significant SNPs annotation.xlsx.

List of *Prakriti* differentiating significant (p<0.05) SNPs in NI and Vadu cohorts.

URL: https://doi.org/10.5281/zenodo.3379895.

**2. Table S2:** Replication NI and Vadu Cohorts.xlsx.

List of *Prakriti* differentiating significant Genes (with or without replicated SNP) & SNPs (with or without replicated profiles) between NI and Vadu cohorts.

URL: https://doi.org/10.5281/zenodo.3381026.

**3. Table-S3:** Replicated 110 SNPs.xlsx.

List of replicated 110 SNPs (with exact profiles) between NI and Vadu cohorts.

URL: https://doi.org/10.5281/zenodo.3381048.

**4. Table S4:** Power Analysis.xlsx.

Power estimation for each significant SNPs from NI and Vadu cohorts.

URL: https://doi.org/10.5281/zenodo.3381054.

**5. Table S5:** Biological Process Summary (shared genes).xlsx.

Compiled data for Gene Ontology analysis done using TOPPFUN for significant genes from pair-wise *Prakriti* comparisons (P *vs* K,V *vs* K and V *vs* P).

URL: https://doi.org/10.5281/zenodo.3402359.

**6. Table S6:** GWAS Exact SNPs.xlsx.

List of *Prakriti* differentiating SNPs from NI and Vadu cohorts associated with disease/traits in GWAS catalog v1.0.2.

URL: https://doi.org/10.5281/zenodo.3383131.

**7. Table S7:** GWAS LD SNPs.xlsx.

List of SNPs associated with disease/traits in GWAS catalog1.0.2 in strong LD (r^2^>0.8) with *Prakriti* differentiating SNPs in NI and Vadu cohorts.

URL: https://doi.org/10.5281/zenodo.3383142.

**8. Table S8:** PheWAS LD SNPs.xlsx.

List of SNPs associated with multiple phenotypes in PheWAS catalog in strong LD (r^2^>0.8) with *Prakriti* differentiating SNPs in NI and Vadu cohorts.

URL: https://doi.org/10.5281/zenodo.3383150.

**9. Table S9:** eQTL’s in *Prakriti* replicated SNPs.xlsx.

List of *Prakriti* replicated SNPs acting as cis-eQTL in GTEx tissues.

URL: https://doi.org/10.5281/zenodo.3402269.

**10. Table S10:** Random Forest Ranks.xlsx.

Details of random forest model results based on (i) genotype to feature and (ii) feature to *Prakriti*.

URL: https://doi.org/10.5281/zenodo.3402273.

**11. Table S11:** Discussion SNPs.xlsx.

Details of replicated SNPs mentioned in discussion section.

URL: https://doi.org/10.5281/zenodo.3402279.

**12. Table S12:** Mouse Knockdown/knockout Phenotypes of Replicated 40 genes.xlsx.

Mouse Knockdown/knockout Phenotypes of Replicated 40 genes.

URL: https://doi.org/10.5281/zenodo.3565519.

## Additional Files 2

### Supplementary figures

1. **Fig. S1:** Distribution of significant SNPs across genomic regions from NI and Vadu cohorts in each pair-wise *Prakriti* comparison. Exonic and 3’UTR have maximum number of differentiating SNPs. The number above the bars indicates the number of differentiating SNPs obtained in each region.
2. **Fig. S2:** Box plot showing estimated power for each genotype with varying sample size in each pair-wise *Prakriti* comparison (A) NI and (B) Vadu cohort.
3. **Fig. S3:** PCA plot of (A) replicated 472 significant SNPs (372 genes) and (B) replicated 110 SNPs (with exact profiles) in NI and Vadu cohort using EGIENSTRAT.
4. **Fig. S4:***ANKLE1* locus that is a pleiotropic modifier locus for breast and ovarian cancer exhibits differences between the *Prakriti* types in both cohorts. (a) Alleles of *ANKLE1* SNPs (rs8100241 and rs8108174) exhibit differences between Vata and Kapha in NI and Vadu cohorts. Vata associated *ANKLE1* SNPs are associated correlated with lower expression of *ABHD8* in GTEx data. (b) Schematic from Figure 3(b) of Lawreson K et. al. (PMID: 27601076) showing the physical map of 13 risk-associated SNPs (red color), the *ABHD8* promoter fragment (green) and the position of the interacting NcoI fragment (purple bar); which was demonstrated by 3C interaction. (c) *ABHD8* knockdown phenotypes in mouse retrieved from MGI database; the phenotypes are associated with anatomical and behavioral phenotypes some of which resonate with *Prakriti* attributes. (d) rs8100241 is in strong LD with PheWAS SNP (rs2363956); bubble plot of PheWAS associations show multiple seemingly unrelated phenotypes associated with rs2363956 of *ANKLE1* gene.
5. **Fig. S5:** Cumulative frequency plots of percentage of significant SNPs in GTEx tissues in pair-wise *Prakriti* comparisons (P*vs*K, V*vs*K and V*vs*P). The x-axis represents the cumulative proportion of differentiating SNPs that are eQTLs across diverse GTEX tissues represented in the y-axis in each *Prakriti* group comparisons across both cohorts.The patterns of cis-eQTL enrichment is similar across both cohorts with significant enrichments in tissues such as testis, nerve tibia, adipose and depletion in uterus, vagina etc.
6. **Fig. S6:** Bubbleplot heatmap representing the eQTL effect size of alternate alleles of SNPs in (A) *IFIT5* and (B) *ZNF502* retrieved from GTEx V7 cis-eQTL data. The effect of *Prakriti* differentiating SNP has been outlined across tissues.
7. **Fig. S7:** Alluvial plot of *FIG4* gene and its connectivity to various biological processes and mouse phenotypes. This highlights that extreme *Prakriti* phenotypes could yield genes with pleiotropic effects from development to physiology.
8. **Fig. S8:** Bubble plot represent the frequency of occurrence of feature values of 17 phenotypes that are significantly associated rs7244213 of *SLC14A2* based on the random forest model.
9. **Fig. S9:** Distinct functions described for extreme *Prakriti* types in Ayurveda.

## References

1. Topol EJ. Individualized medicine from prewomb to tomb. Cell. 2014;157:241–253.

2. Snyder M, Weissman S, Gerstein M. Personal phenotypes to go with personal genomes. Molecular systems biology. 2009;5:273.

3. Loscalzo J, Barabasi A-L. Systems biology and the future of medicine. Wiley interdisciplinary reviews Systems biology and medicine. 2011;3:619–627.

4. Chen R, Snyder M. Promise of personalized omics to precision medicine. Wiley interdisciplinary reviews Systems biology and medicine. 2013;5:73–82.

5. Price ND, Magis AT, Earls JC, Glusman G, Levy R, Lausted C, et al. A wellness study of 108 individuals using personal, dense, dynamic data clouds. Nat Biotechnol. 2017;35:747–56.

6. Tam V, Patel N, Turcotte M, Bossé Y, Paré G, Meyre D. Benefits and limitations of genome-wide association studies. Nature Reviews Genetics. 2019;1.

7. Goh K-I, Cusick ME, Valle D, Childs B, Vidal M, Barabási A-L. The human disease network. Proceedings of the National Academy of Sciences of the United States of America. 2007;104:8685–8690.

8. Welter D, MacArthur J, Morales J, Burdett T, Hall P, Junkins H, et al. The NHGRI GWAS Catalog, a curated resource of SNP-trait associations. Nucleic acids research. 2014;42:D1001–D1006.

9. Bush WS, Oetjens MT, Crawford DC. Unravelling the human genome-phenome relationship using phenome-wide association studies. Nature reviews Genetics. 2016;17:129–145.

10. Denny JC, Bastarache L, Ritchie MD, Carroll RJ, Zink R, Mosley JD, et al. Systematic comparison of phenome-wide association study of electronic medical record data and genome-wide association study data. Nat Biotechnol. 2013;31:1102–10.

11. Millard LAC, Davies NM, Tilling K, Gaunt TR, Davey Smith G. Searching for the causal effects of body mass index in over 300 000 participants in UK Biobank, using Mendelian randomization. PLoS genetics. 2019;15:e1007951.

12. Verma A, Lucas A, Verma SS, Zhang Y, Josyula N, Khan A, et al. PheWAS and Beyond: The Landscape of Associations with Medical Diagnoses and Clinical Measures across 38,662 Individuals from Geisinger. American journal of human genetics. 2018;102:592–608.

13. Pendergrass SA, Brown-Gentry K, Dudek S, Frase A, Torstenson ES, Goodloe R, et al. Phenome-wide association study (PheWAS) for detection of pleiotropy within the Population Architecture using Genomics and Epidemiology (PAGE) Network. PLoS genetics. 2013;9:e1003087.

14. Cai T, Zhang Y, Ho Y-L, Link N, Sun J, Huang J, et al. Association of Interleukin 6 Receptor Variant With Cardiovascular Disease Effects of Interleukin 6 Receptor Blocking Therapy: A Phenome-Wide Association Study. JAMA Cardiol. 2018;3:849–57.

15. Rastegar-Mojarad M, Ye Z, Kolesar JM, Hebbring SJ, Lin SM. Opportunities for drug repositioning from phenome-wide association studies. Nature Biotechnology. 2015;33:342–5.

16. Diogo D, Tian C, Franklin CS, Alanne-Kinnunen M, March M, Spencer CCA, et al. Phenome-wide association studies across large population cohorts support drug target validation. Nature communications. 2018;9:4285.

17. Yin W, Gao C, Xu Y, Li B, Ruderfer DM, Chen Y. Learning Opportunities for Drug Repositioning via GWAS and PheWAS Findings. AMIA Joint Summits on Translational Science proceedings AMIA Joint Summits on Translational Science. 2018;2017:237–246.

18. Hebbring SJ. The challenges, advantages and future of phenome-wide association studies. Immunology. 2014;141:157–65.

19. Ried JS, M JJ, Chu AY, Bragg-Gresham JL, van Dongen J, Huffman JE, et al. A principal component meta-analysis on multiple anthropometric traits identifies novel loci for body shape. Nature Communications. 2016;7:1–11.

20. Emond MJ, Louie T, Emerson J, Zhao W, Mathias RA, Knowles MR, et al. Exome sequencing of extreme phenotypes identifies DCTN4 as a modifier of chronic Pseudomonas aeruginosa infection in cystic fibrosis. Nature genetics. 2012;44:886–889.

21. Emond MJ, Louie T, Emerson J, Chong JX, Mathias RA, Knowles MR, et al. Exome Sequencing of Phenotypic Extremes Identifies CAV2 and TMC6 as Interacting Modifiers of Chronic Pseudomonas aeruginosa Infection in Cystic Fibrosis. PLoS genetics. 2015;11:e1005273.

22. Aubart M, Gazal S, Arnaud P, Benarroch L, Gross M-S, Buratti J, et al. Association of modifiers and other genetic factors explain Marfan syndrome clinical variability. European journal of human genetics: EJHG. 2018;26:1759–1772.

23. Fay AP, de Velasco G, Ho TH, Van Allen EM, Murray B, Albiges L, et al. Whole-Exome Sequencing in Two Extreme Phenotypes of Response to VEGF-Targeted Therapies in Patients With Metastatic Clear Cell Renal Cell Carcinoma. J Natl Compr Canc Netw. 2016;14:820–4.

24. Chan Y, Holmen OL, Dauber A, Vatten L, Havulinna AS, Skorpen F, et al. Common variants show predicted polygenic effects on height in the tails of the distribution, except in extremely short individuals. PLoS Genet. 2011;7:e1002439.

25. Sethi TP, Prasher B, Mukerji M. Ayurgenomics: a new way of threading molecular variability for stratified medicine. ACS Chem Biol. 2011;6:875–80.

26. Sagner M, McNeil A, Puska P, Auffray C, Price ND, Hood L, et al. The P4 Health Spectrum – A Predictive, Preventive, Personalized and Participatory Continuum for Promoting Healthspan. Progress in Cardiovascular Diseases. 2017;59:506–21.

27. Prasher B, Gibson G, Mukerji M. Genomic insights into ayurvedic and western approaches to personalized medicine. J Genet. 2016;95:209–28.

28. Prasher B, Negi S, Aggarwal S, Mandal AK, Sethi TP, Deshmukh SR, et al. Whole genome expression and biochemical correlates of extreme constitutional types defined in Ayurveda. J Transl Med. 2008;6:48.

29. Tiwari P, Kutum R, Sethi T, Shrivastava A, Girase B, Aggarwal S, et al. Recapitulation of Ayurveda constitution types by machine learning of phenotypic traits. PLoS ONE. 2017;12:e0185380.

30. Govindaraj P, Nizamuddin S, Sharath A, Jyothi V, Rotti H, Raval R, et al. Genome-wide analysis correlates Ayurveda Prakriti. Scientific reports. 2015;5:15786.

31. Rotti H, Mallya S, Kabekkodu SP, Chakrabarty S, Bhale S, Bharadwaj R, et al. DNA methylation analysis of phenotype specific stratified Indian population. Journal of translational medicine. 2015;13:151.

32. Aggarwal S, Negi S, Jha P, Singh PK, Stobdan T, Pasha MAQ, et al. EGLN1 involvement in high-altitude adaptation revealed through genetic analysis of extreme constitution types defined in Ayurveda. Proc Natl Acad Sci USA. 2010;107:18961–6.

33. Chauhan NS, Pandey R, Mondal AK, Gupta S, Verma MK, Jain S, et al. Western Indian Rural Gut Microbial Diversity in Extreme Prakriti Endo-Phenotypes Reveals Signature Microbes. Front Microbiol. 2018;9:118.

34. Aggarwal S, Gheware A, Agrawal A, Ghosh S, Prasher B, Mukerji M, et al. Combined genetic effects of EGLN1 and VWF modulate thrombotic outcome in hypoxia revealed by Ayurgenomics approach. J Transl Med. 2015;13:184.

35. Prasher B, Varma B, Kumar A, Khuntia BK, Pandey R, Narang A, et al. Ayurgenomics for stratified medicine: TRISUTRA consortium initiative across ethnically and geographically diverse Indian populations. J Ethnopharmacol. 2017;197:274–93.

36. Sasayama D, Hattori K, Ogawa S, Yokota Y, Matsumura R, Teraishi T, et al. Genome-wide quantitative trait loci mapping of the human cerebrospinal fluid proteome. Human molecular genetics. 2017;26:44–51.

37. Ahola-Olli AV, Würtz P, Havulinna AS, Aalto K, Pitkänen N, Lehtimäki T, et al. Genome-wide Association Study Identifies 27 Loci Influencing Concentrations of Circulating Cytokines and Growth Factors. American journal of human genetics. 2017;100:40–50.

38. Kitsios GD, Tangri N, Castaldi PJ, Ioannidis JPA. Laboratory mouse models for the human genome-wide associations. PLoS ONE. 2010;5:e13782.

39. Consortium Gte, Laboratory DA &Coordinating C (LDACC)—Analysis WG, Group SM groups-AW, groups EGte (eGTEx), Fund NC, NIH/NCI, et al. Genetic effects on gene expression across human tissues. Nature. 2017;550:204–213.

40. Bult CJ, Eppig JT, Blake JA, Kadin JA, Richardson JE, Group MGD. The mouse genome database: genotypes, phenotypes, and models of human disease. Nucleic acids research. 2013;41:D885–D891.

41. Lawrenson K, Kar S, McCue K, Kuchenbaeker K, Michailidou K, Tyrer J, et al. Functional mechanisms underlying pleiotropic risk alleles at the 19p13.1 breast–ovarian cancer susceptibility locus. Nature Communications. 2016;7:12675.

42. Kleinstein SE, Rein M, Abdelmalek MF, Guy CD, Goldstein DB, Mae Diehl A, et al. Whole-Exome Sequencing Study of Extreme Phenotypes of NAFLD. Hepatology communications. 2018;2:1021–1029.

43. Ferreira RC, Freitag DF, Cutler AJ, Howson JMM, Rainbow DB, Smyth DJ, et al. Functional IL6R 358Ala allele impairs classical IL-6 receptor signaling and influences risk of diverse inflammatory diseases. PLoS Genet. 2013;9:e1003444.

44. Vargas VRA, Bonatto SL, Macagnan FE, Feoli AMP, Alho CS, Santos NDV, et al. Influence of the 48867A>C (Asp358Ala) IL6R polymorphism on response to a lifestyle modification intervention in individuals with metabolic syndrome. Genet Mol Res. 2013;12:3983–91.

45. Leo B. Random forests. Machine learning. 2001;45:5–32.

46. Kipper S, Hamad S, Caly L, Avrahami D, Bacharach E, Jans DA, et al. New host factors important for respiratory syncytial virus (RSV) replication revealed by a novel microfluidics screen for interactors of matrix (M) protein. Mol Cell Proteomics. 2015;14:532–43.

47. Liu J, Shen L, Yang L, Hu S, Xu L, Wu S. High expression of B3GNT8 is associated with the metastatic potential of human glioma. Int J Mol Med. 2014;33:1459–68.

48. Juyal RC, Negi S, Wakhode P, Bhat S, Bhat B, Thelma BK. Potential of ayurgenomics approach in complex trait research: leads from a pilot study on rheumatoid arthritis. PLoS ONE. 2012;7:e45752.

49. Knepper MA, Kwon T-H, Nielsen S. Molecular physiology of water balance. The New England journal of medicine. 2015;372:1349–1358.

50. Xu S, Yang Y, Zhou X, Xu J, Zhou K, Yang G. Adaptive evolution of the osmoregulation-related genes in cetaceans during secondary aquatic adaptation. BMC Evol Biol. 2013;13:189.

51. Hong X, Xing H, Yu Y, Wen Y, Zhang Y, Zhang S, et al. Genetic polymorphisms of the urea transporter gene are associated with antihypertensive response to nifedipine GITS. Methods Find Exp Clin Pharmacol. 2007;29:3–10.

52. Knepper MA, Miranda CA. Urea channel inhibitors: a new functional class of aquaretics. Kidney Int. 2013;83:991–3.

53. Consortium IGV. The Indian Genome Variation database (IGVdb): a project overview. Human genetics. 2005;118:1–11.

54. Wang K, Li M, Hakonarson H. ANNOVAR: functional annotation of genetic variants from high-throughput sequencing data. Nucleic Acids Res. 2010;38:e164.55.

55. Giner G, Smyth GK. statmod: probability calculations for the inverse Gaussian distribution. R Journal. 2016;8:339–51.

56. Price AL, Patterson NJ, Plenge RM, Weinblatt ME, Shadick NA, Reich D. Principal components analysis corrects for stratification in genome-wide association studies. Nat Genet. 2006;38:904–9.

57. Chen J, Bardes EE, Aronow BJ, Jegga AG. ToppGene Suite for gene list enrichment analysis and candidate gene prioritization. Nucleic Acids Res. 2009;37:W305–311.

58. Arnold M, Raffler J, Pfeufer A, Suhre K, Kastenmüller G. SNiPA: an interactive, genetic variant-centered annotation browser. Bioinformatics. 2015;31:1334–6.

59. Bojanowski M, Edwards R. alluvial: R Package for Creating Alluvial Diagrams [Internet]. 2016. Available from: https://github.com/mbojan/alluvial

60. Kursa MB, Rudnicki WR. Feature Selection with the Boruta Package. Journal of Statistical Software. 2010;36:1–13.

