## Additional Files 2 for "Genetic differences between extreme and composite constitution types from whole exome sequences reveal actionable variations"

**Supplementary Figure 1**

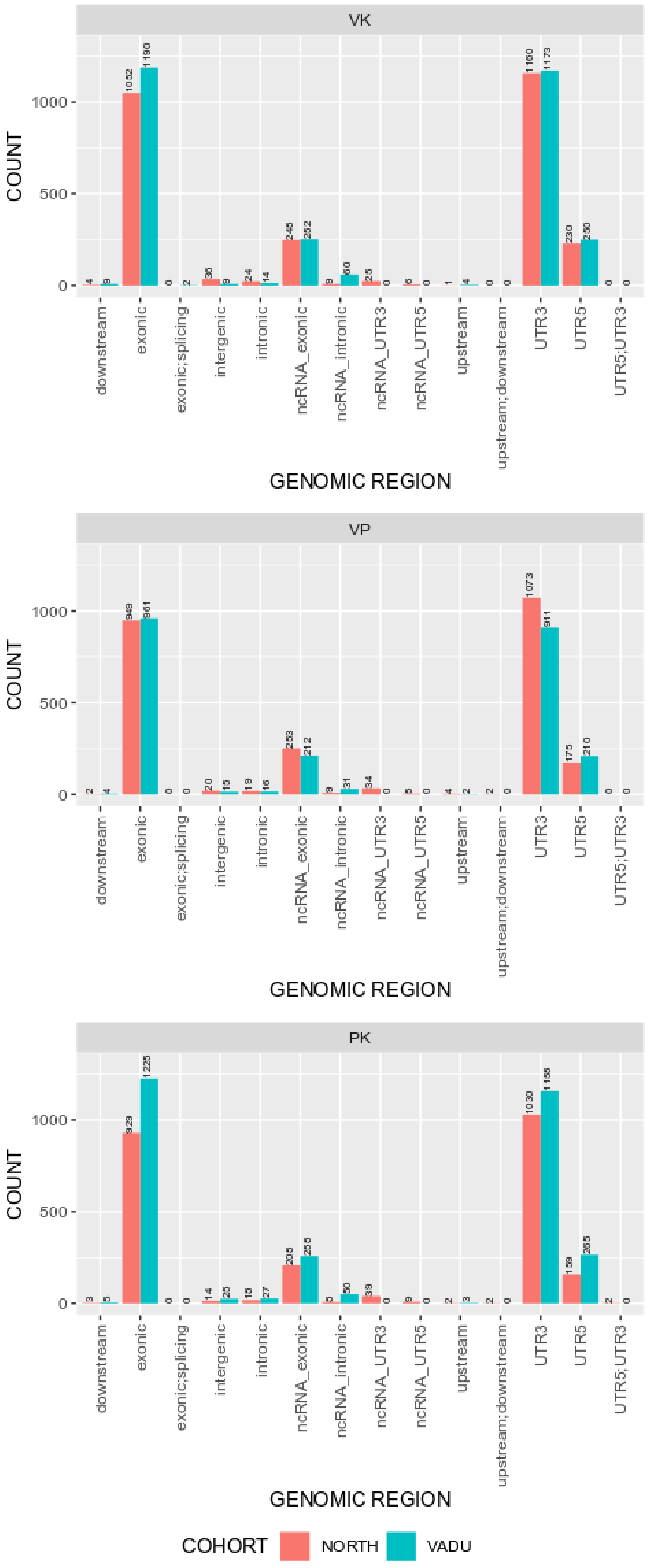

**Supplementary Figure 2**

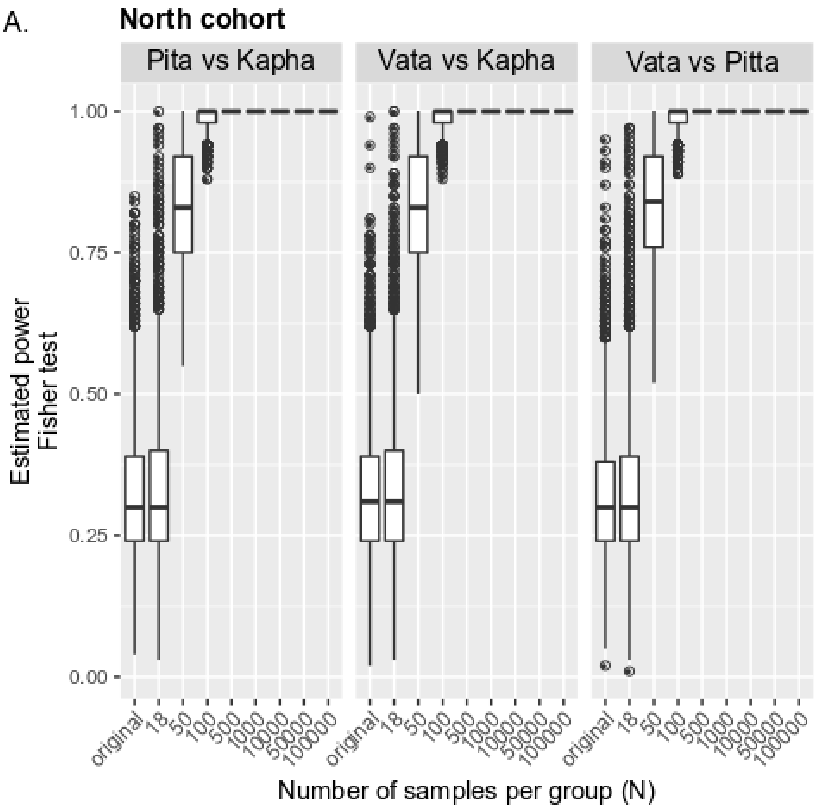


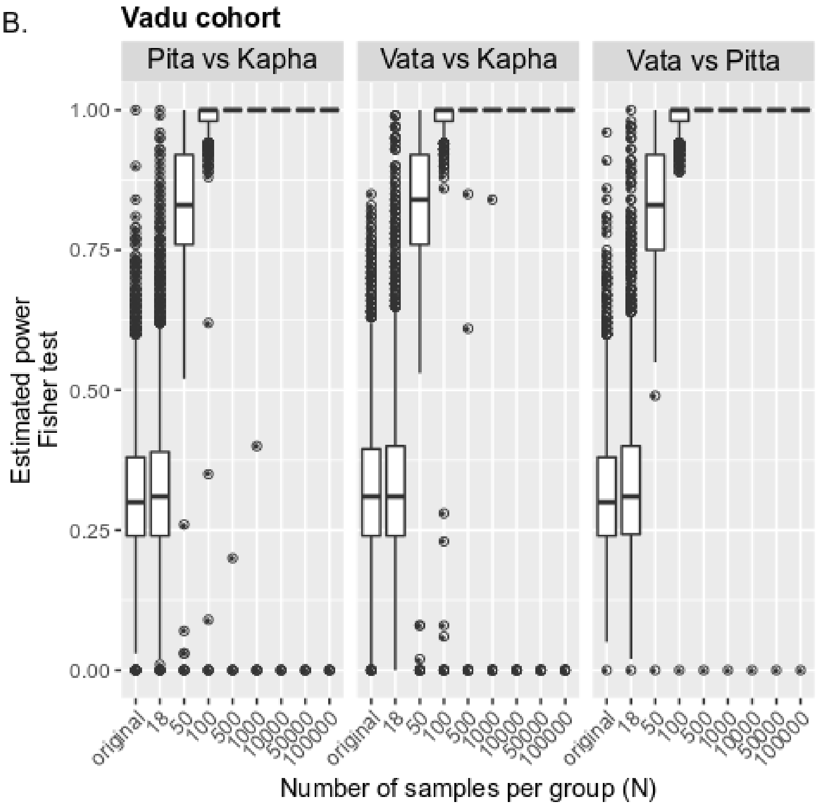

**Supplementary Figure 3**

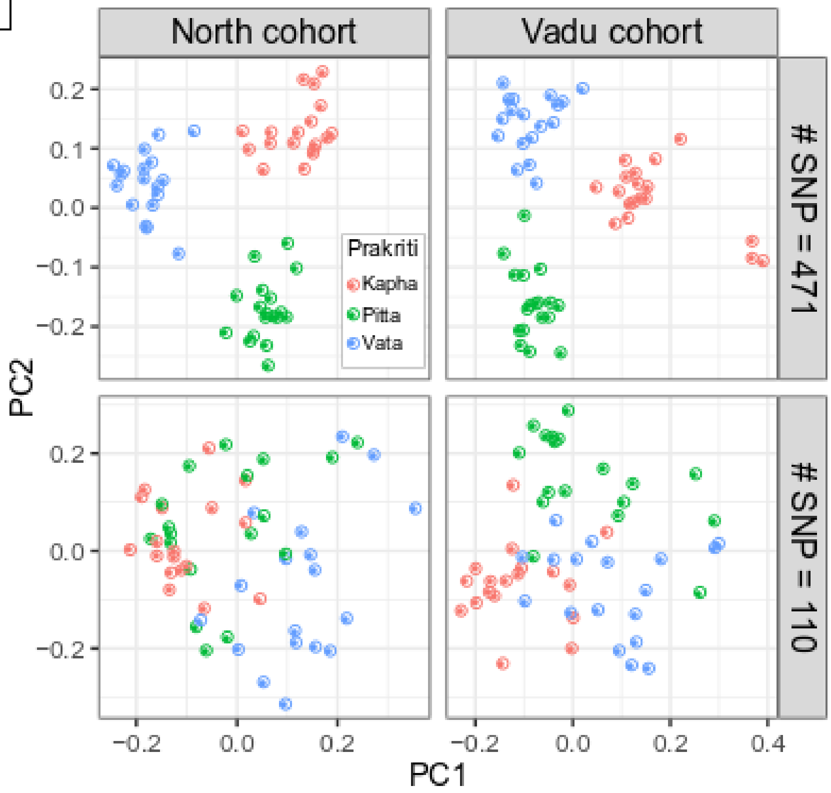

**Supplementary Figure 4**

A.

| dbSNP | Nature | Vata | Kapha | Cohort |
|---|---|---|---|---|
| rs8100241 (ANKLE1) | missense (A to L) | 0.611 | 0.333 | North |
| " | " | 0.611 | 0.294 | Vadu |
| rs8108174 (ANKLE1) | missense (L to Q) | 0.611 | 0.333 | North |
| " | " | 0.583 | 0.277 | Vadu |

B.

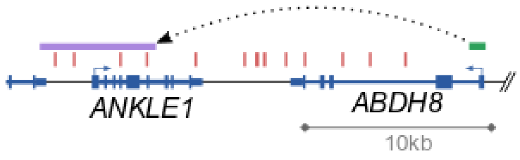


D.

**Mouse knock down phenotype**

- Decrease body length
- Decrease bone mineral content
- Decrease bone mineral density
- Short tibia
- Decreased lean body mass
- Hyperactivity
- Increased exploration in new environment

D.

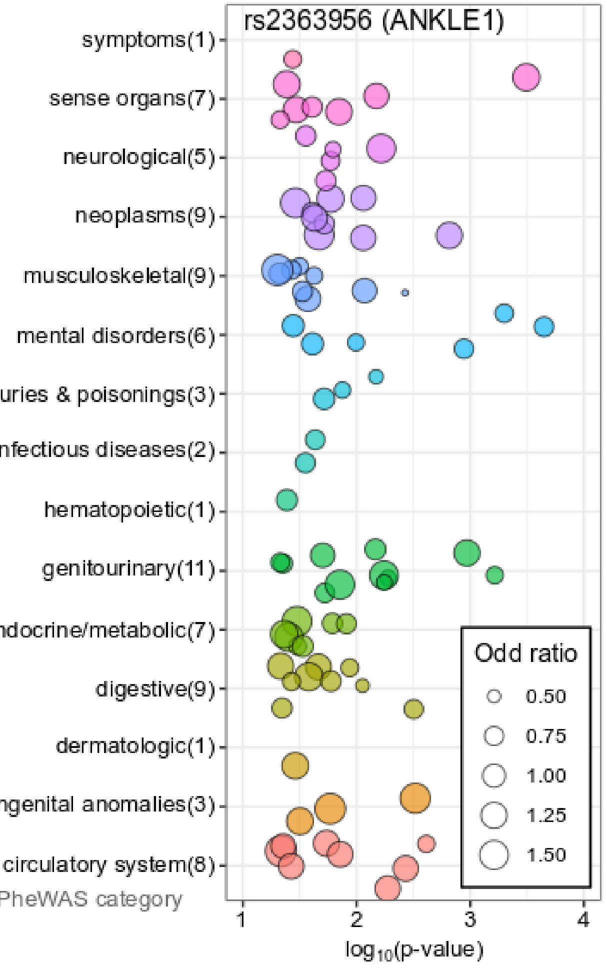

**Supplementary Figure 5**

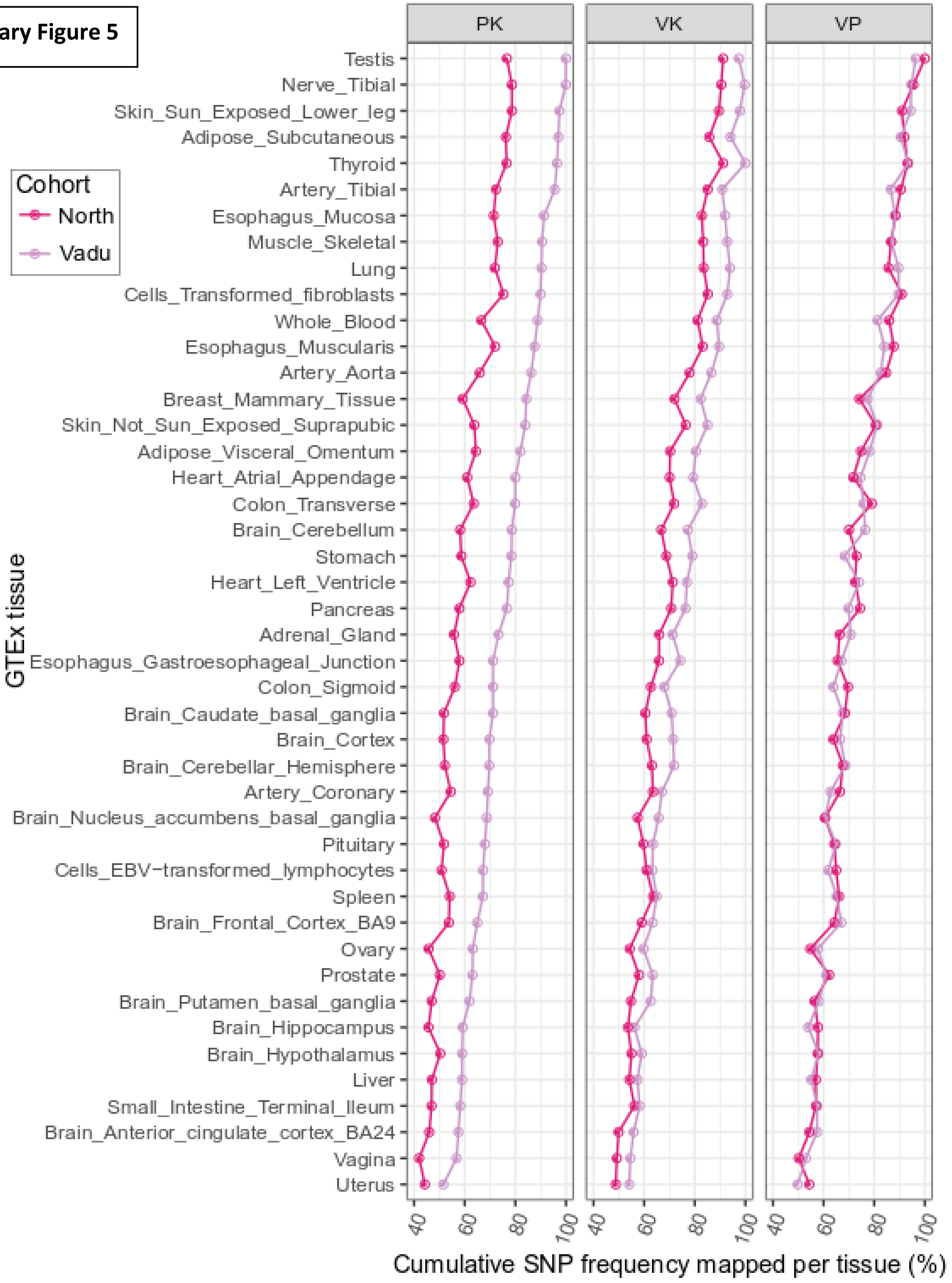

**Supplementary Figure 6**

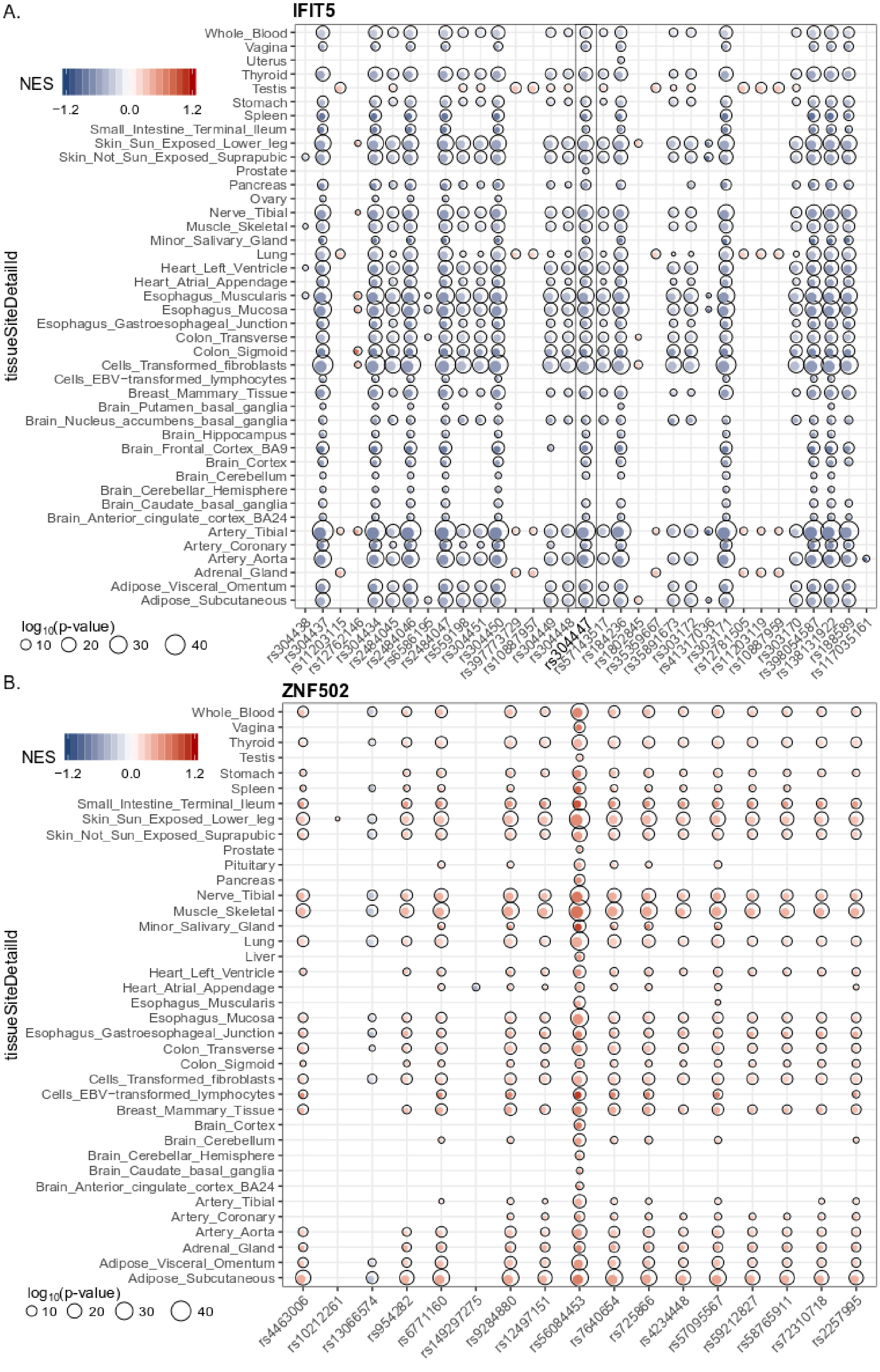

**Supplementary Figure 7**

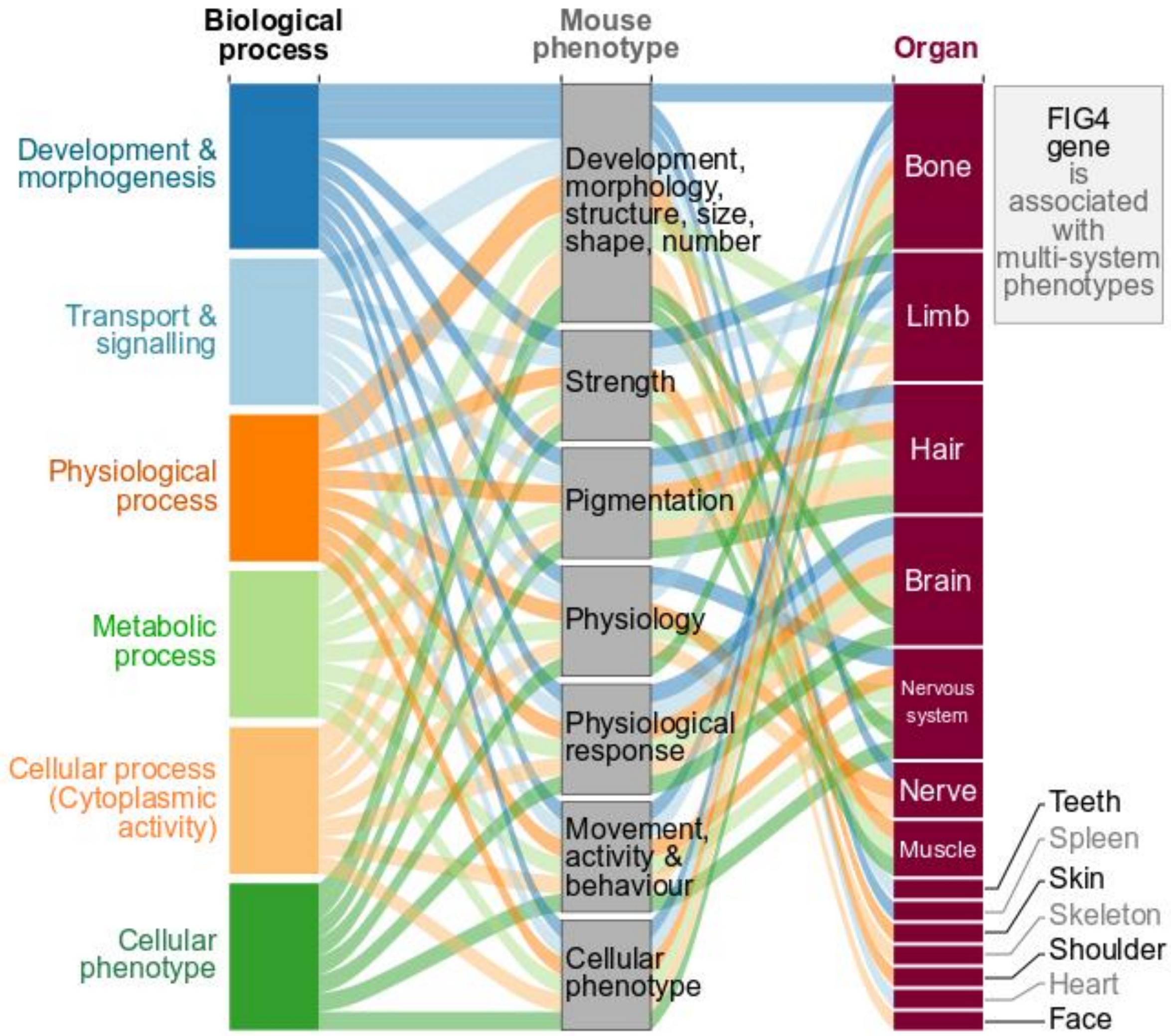

**Supplementary Figure 8**

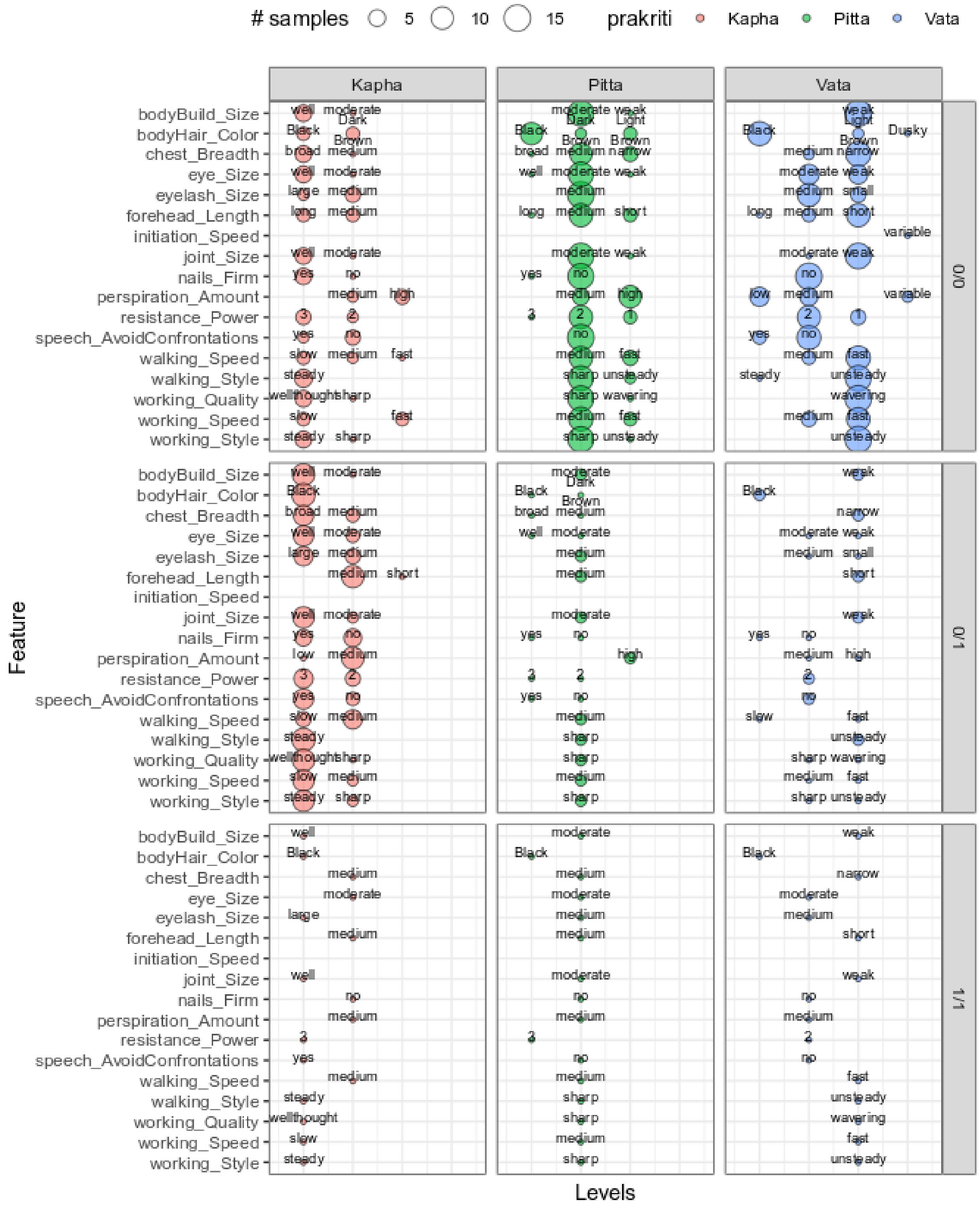

**Supplementary Figure 9**

### Distinct functions have been ascribed for *Vata*, *Pitta* and *Kapha*

***Vata* functions:**

Pulsatile action, transport, absorption, sorting, excretion, Through Maintenance and control of movements and activity

प्रस्पन्दनोद्वहनपूरणविवेकधारणलक्षणो वायुः पञ्चधा प्रविभक्तः शरीरं धारयति

[illegible]

[illegible] ||४९||

***Pitta* functions:**

Digestion, metabolism, Pigmentation, skin coloration, immuno-metabolism, thought- intellect, Through maintenance of metabolic balance

रागपक्त्योजस्तेजोमेधोष्मकृत् पित्तं पञ्चधा प्रविभक्तमग्निकर्मणाऽनुग्रहं करोति

[illegible]

[illegible] ||५०||

***Kapha* functions:**

Lubrication of joints, skin, and body in general, storage, Bulk formation, healing, strength & stability through maintenance of Water Balance

सन्धिसंश्लेषणस्नेहनरोपणपूरणबलस्थैर्यकृच्छ्लेष्मा पञ्चधा प्रविभक्त

उदककर्मणाऽनुग्रहं करोति ||४||

[illegible]

[illegible] ||५१||  -C. Su. 18/49-51
